# A theoretical and experimental framework enables low-coverage sequencing for accurate quantification of genome-wide cytosine modification levels

**DOI:** 10.1101/2025.01.08.631958

**Authors:** Christian E. Loo, Johanna M. Fowler, Heqiao Zhu, Christopher Krapp, Ruiyao Zhu, Marisa S. Bartolomei, Wanding Zhou, Rahul M. Kohli

**Author notes:** Correspondence to: Wanding Zhou, Rahul M. Kohli.

## Abstract

5-methylcytosine (5mC) and 5-hydroxymethylcytosine (5hmC) regulate gene expression and exhibit dynamic levels during development and disease. While high-depth, base-resolution studies offer the most detailed view of epigenetic landscapes, many open questions are answered by surveying changes in 5mC/5hmC levels across larger cohorts. Nonetheless, current global quantification methods, including mass spectrometry, are typically limited in accessibility, accuracy, or throughput. Here, to evaluate the viability of low-coverage sequencing as an alternative, we first computationally downsampled deeply sequenced data to resolve the three-way relationship between sequencing coverage, modification levels, and measurement error. This relationship allowed us to develop a facile online tool for error calculation and to define experimental targets: <0.24% genome coverage can quantify 5mC and low-abundance 5hmC with minimal and predictable errors (<5%). Importantly, in direct comparisons, low-depth sequencing (Sparse-Seq) demonstrated high accuracy and less variability than mass spectrometry, while distinctively preserving genomic context. Applied serially to developing mouse brains, Sparse-Seq revealed an earlier emergence of 5hmCpG compared to 5mCpH and uncovered previously overlooked, genomic feature-specific epigenetic changes. This work establishes a rigorous foundation for employing Sparse-Seq as a highly accessible approach for 5mC/5hmC quantification, enabling economical first-pass analysis of epigenetic landscapes suited for large cohort studies and new hypothesis generation.

## INTRODUCTION

5-methylcytosine (5mC) is a critical epigenetic DNA modification in mammalian genomes, most commonly found in cytosine-guanine (CpG) dinucleotide contexts. 5mC is known to typically act as a repressive mark, with key roles in development and differentiation, imprinting, X-chromosome inactivation, and the suppression of mobile genetic elements [1]. The discovery of the ten-eleven translocation (TET) family of enzymes has triggered the more recent focus beyond 5mC to include 5-hydroxymethylcytosine (5hmC), a prevalent and previously underappreciated mark in many cell types [2, 3]. Oxidation of 5mC by TET enzymes initiates the process of DNA demethylation, whereby the repressive 5mC marks are erased [4, 5]. At the same time, the high level and stability of 5hmC in certain cell types have suggested that this epigenetic modification also has its own regulatory roles, which can oppose those of 5mC in many regards [6–8].

In the study of the biological roles for 5mC and 5hmC, significant insights have been gained by localizing and quantifying these DNA modifications [9, 10]. Base-resolution sequencing approaches can be crucial for distinguishing the specific roles of 5mC and 5hmC in individual biological samples. However, these methods can be prohibitively expensive when applied at genome-wide scales, especially when analyzing a large number of biologically important samples. These issues are even more acute when evaluating 5hmC, as its distribution does not follow the bimodal pattern of 5mC and its overall lower abundance requires even higher sequencing depth. In these cases, profiling the changes in global (genome-wide) levels of 5mC and 5hmC has been regarded as a more accessible approach to studying DNA modification dynamics in disease and development. For example, studying genome-wide levels of 5mC and 5hmC has helped show roles for TET enzymes in both early zygotic reprogramming and primordial germ cell development [11, 12]. In post-mitotic tissues such as the brain, levels of 5hmC track with development, reaching levels as high as 20% of those of 5mC, with demonstrated roles in learning and memory [7, 13, 14]. Studying larger cohorts suggests that global 5mC or 5hmC levels can serve as biomarkers in cancer detection or prognosis [15–17], and aberrant global methylation or hydroxymethylation has also been shown in non-cancerous processes, including stroke and neurodegenerative diseases [18–20].

Given the biological significance of 5mC and 5hmC, it is essential to have accurate and sensitive methods for quantification of these cytosine modifications in genomic DNA (gDNA) samples (**Figure 1, left**). While several approaches for measuring cytosine modification levels are available, the most commonly employed involve immunochemistry, mass spectrometry, or sequencing-based techniques (**Figure 1, right)**. Immunochemistry-based techniques such as ELISA with antibodies against 5mC and 5hmC can be facile to implement. However, they are less quantitative than liquid chromatography-tandem mass spectrometry (LC-MS/MS), which has been generally regarded as the ‘gold standard’ [21–23]. With LC-MS/MS, gDNA is purified and degraded to nucleosides. The nucleosides are then separated by LC, identified by MS/MS, and quantified by comparison to standards. Although this approach is highly specific and sensitive, its accessibility is constrained by the relatively high input amounts of DNA required (typically >500 ng), the need for rigorous protocol optimization, and the necessity of instrument availability [24–26]. Iterations to LC-MS/MS methods have sought to reduce the input requirements; however, several limitations remain. Most notably, due to the obligate degradation of DNA to individual nucleosides prior to analysis, LC-MS/MS cannot report the genomic locations or sequence context of these modifications. This is a significant limitation, given the distinct biological implications for DNA modifications in CpG versus non-CpG contexts or in different genomic elements.

**Figure 1.**
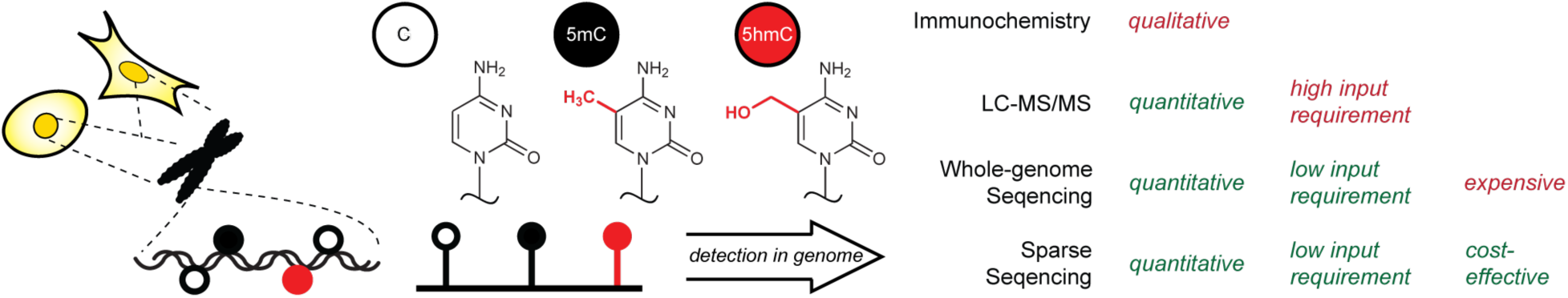
Methods for detecting cytosine modifications. Structures of the nucleobase cytosine and its two most common mammalian genomic variants, 5-methylcytosine and 5-hydroxymethylcytosine (left). Common methods for quantifying cytosine modifications in the genome and their potential strengths and limitations are noted (right).

Sequencing of gDNA samples offers a common alternative for measuring global DNA modification levels. Most efforts center around bisulfite sequencing (BS-Seq), whereby unmodified cytosines are chemically deaminated by sodium bisulfite and next-generation sequencing (NGS) is used to resolve unmodified and modified bases at base resolution [27]. Conventional BS-Seq itself is unable to resolve 5mC and 5hmC, although methods coupling bisulfite to enzymatic oxidation (TAB-Seq), chemical oxidation (oxBS-Seq), or enzymatic deamination (bACE-Seq) have been developed to distinguish these two most prevalent modifications [28–31]. Non-destructive alternatives to bisulfite-based approaches have also been recently developed, including all-enzymatic methods for profiling cytosine modifications such as enzymatic-methylation sequencing (EM-Seq, 5mC + 5hmC), APOBEC-coupled epigenetic sequencing (ACE-Seq, 5hmC alone), Direct Methylation sequencing (DM-Seq, 5mC alone), or hybrid enzymatic-chemical methods such as TET-assisted pyridine borane sequencing (TAPS) [32–35].

The most rigorous application of sequencing involves whole-genome base-resolution analysis. However, such an approach can be cost-prohibitive, especially when base-resolution information is not absolutely required. Modifications to these protocols have, therefore, been employed to estimate global cytosine modification levels. Such approaches include repetitive element pyrosequencing, reduced representation bisulfite sequencing (RRBS), or immunoenrichment (MeDIP), although each approach brings different potential sets of bias to sampling of the genome [36–38]. Similarly, analysis of gDNA with array-based assays can provide coverage of a large number of specific genomic sites. However, the array probes are biased to query some genomic elements more than others [39], and the validity of such approaches in probing global 5hmC levels is not well-established.

Recently, low-coverage BS-Seq has been suggested as a plausible alternative to deep whole-genome BS-Seq (WGBS) for estimating global DNA modification levels. Theoretical estimates led to the proposal that sequencing as few as 10,000 total cytosines in a CpG context could provide minimal margins of error for global levels of aggregate cytosine modifications [40]. Motivated by these calculations, we aimed to explore the potential generalizability of low-depth sequencing, which we term as Sparse Sequencing (Sparse-Seq), as a facile, economical, and accurate method for estimating genome-wide levels of both 5mC and 5hmC. While shallow sequencing is often employed by users to pre-screen libraries prior to deep sequencing, we recognized that no prior framework has been established for understanding the tradeoffs between accuracy and sequencing depth, nor for calculating the error associated with given measurements derived from low-coverage sequencing. This critical gap prevents low-coverage sequencing from being deployed as a rigorous quantitative method for profiling 5mC and 5hmC as there is no basis for knowing when a measurement is trustworthy or how to compare estimates across samples. Here, using computational analysis, we first demonstrate that sparse sampling can theoretically be applied with high accuracy, even when modifications such as 5hmC drop to low levels. We then translate this robust theoretical analysis into an error calculation tool that empowers users to make principled, confidence-aware sequencing decisions. Guided by this tool, we then experimentally benchmark Sparse-Seq against LC-MS/MS, with a specific emphasis on a combined chemical and enzymatic (bACE-Seq) approach that allows for separate resolution of 5mC and 5hmC. Our work demonstrates that Sparse-Seq offers comparable or superior fidelity when compared to LC-MS/MS while offering additional advantages. Unlike LC-MS/MS, Sparse-Seq retains sequence context and can parse changes in modification levels at specific genomic elements, such as enhancers or promoters, rather than just providing global measurements. We therefore offer both strong conceptual validation and the needed tools to utilize low-coverage sequencing as means to accurate and economical profiling of global changes in both methylation and hydroxymethylation in biological samples of interest.

## MATERIAL AND METHODS

### Computational processing and downsampling

ACE-seq, TAB-seq, and EM-seq datasets from murine brain, mouse ESCs, and human cell lines were acquired through GEO using the following accessions: GSE48519, GSE63137, GSE116016, GSE47966, GSE195752. FASTQ reads were trimmed to remove sequencing adapters and aligned to the reference genome assembly (GRCh38 for human and GRCm38 for mouse). Duplicate reads were masked using Picard tools (https://broadinstitute.github.io/picard/). DNA modification levels were extracted using bismark_methylation_extractor. Double counting from the overlapping mate reads was excluded.

To mimic real sparse sequencing data, the deep sequencing datasets were downsampled at the BAM file level to read counts evenly spaced on a log_2_ scale from N= 64 (2^6^) to N= 262,144 (2^18^) reads. The sampled read numbers were then transformed to sequencing depth based on read length and genome size. Each downsampled BAM file went through the same preprocessing steps including duplicate marking and methylation extraction. The cytosine modification levels at CpG, CpHpH, and CpHpG were aggregated to allow for model validation and construction independent of sequence context.

### Estimation of Chromatin State Proportions

To estimate the proportion of each chromatin state in the above public datasets, processed and aligned BAM files were overlaid with a BED file annotated with 18 distinct chromatin states using BEDtools intersect. Chromatin states were defined based on a consensus generated from chromatin state profiles across 66 mouse tissues [41]. For each dataset, the number of reads overlapping each chromatin state was calculated. These counts were then normalized by dividing by the total number of reads in the corresponding dataset to obtain the proportional representation of each chromatin state.

### Downsampling for Chromatin States Analysis

The public datasets were downsampled at the BAM file level to read counts evenly spaced on a log_2_ scale from N= 8,192 (2^6^) to N= 33,554,432 (2^25^) reads. For each downsampled simulation, the cytosine modification levels at CpG were classified into chromatin states and then aggregated using the command line tool YAME. The source of chromatin states was defined as above.

### Interpolation and construction of TAE estimator

A total of 18 datasets (5 TAB-seq, 5 BS-seq, 4 ACE-seq and 4 EM-seq) were combined. Using this data set, LOESS regression (R stats package, degree = 1, span = 0.75) of the total analytical error was performed on log_2_-transformed genome coverage and cytosine modification levels. The regression model was then used to interpolate the TAE for genome coverage input ranging from 0.001% to 1% and total cytosine modification from 0% to 5%. The TAE estimator was constructed to allow for automated conversions of typical input data (https://zhou-lab.github.io/TAE_calculator/). For a given low-coverage sequencing sample, the user performs read mapping and calculation of the modification level of interest (beta value) in either all mapped reads or in a specific ChromHMM genomic element of interest. The modifications of interest can be unmodified C, 5mC, or 5hmC and can be quantified across all C, CpG only, or CpH only. The estimator converts the input modification beta value into a % of total C, adjusting for CpG or CpH context as needed. It additionally calculates genomic coverage based on the input number of reads, read length, and adjusts for desired ChromHMM genomic elements. The interpolated TAE model is then used to output TAE percent and provide the resulting beta value ± error.

### Mouse brain development genomic DNA extraction

Breeding stocks of C57BL/6J (The Jackson Laboratory, Bar Harbor, ME, USA) were maintained in a pathogen-free facility in a 12-hour light-dark cycle. All animals were housed in polysulfone cages and had access to drinking water and chow (Laboratory Autoclavable Rodent Diet 5010, LabDiet) *ad libitum*. Neonatal (P0) brains and postnatal cortices were isolated from mice at indicated ages and snap frozen. Frontal cortices were obtained by slicing the adult brains coronally at Bregma 1 mm, and dissecting the frontal cortical tissue, being careful to avoid contamination with olfactory or putamen regions, under a dissection microscope. All animal work was conducted with the approval of the Institutional Animal Care and Use Committee (IACUC) at the University of Pennsylvania.

To isolate gDNA, brain was thawed and digested in lysis buffer (50 mM Tris, pH 8.0, 100 mM EDTA, 0.5% SDS) with proteinase K (Sigma-Aldrich) overnight at 55°C. Genomic DNA was isolated using Phenol:Chloroform:Isoamyl Alcohol (Fisher BP17501-400) and ethanol precipitation. DNA samples resuspended in TE were stored at 4°C.

## LC-MS/MS

The purified gDNA samples (250 ng) were digested using Nucleotide Digestion Mix (NEB) in a volume of 20 µL at 37°C for 16 hrs to obtain single nucleosides. To 9 µL of the incubated digestion mix, 1 µL of 1% formic acid was added to yield a final solution of 0.1% formic acid. 5 µL of the sample or standard was injected into an HPLC-QQQ-MS/MS system. An Accucore Vanquish C18+ ultra-high performance liquid chromatography system (Thermo, 1.5 µm particle size, 50 mm length) was used to separate nucleosides with online mass spectrometry detection by the Altis (Thermo Fisher Scientific) triple-quadrupole LC mass spectrometer. Nucleosides were separated starting with a 2 minute flow of 100% Buffer A (0.1% formic acid) followed by a 10 minute gradient of 0-4% Buffer B (30% acetonitrile, 0.1% formic acid) at 0.4 mL/min. Mass transitions for deoxynucleosides were C 228.093→112.05 *m/z*; 5mC 242.108→126.066 *m/z*; and 5hmC 258.103→142.061 *m/z*. Standard curves were generated using standard nucleosides ranging from 5.56 fmol to 92500 fmol for unmodified cytosine and 0.565 fmol to 9250 fmol for modified cytosines.

### BS/bACE-Seq library prep

Purified mouse gDNA (200 ng) with 1 ng each of phiX, mCpG pUC19 (Zymo), and 5hmC T4 spike-in controls were sheared to approximately 350 bp using a Covaris instrument. Sheared gDNA was taken through end prep and adaptor ligation using the NEBNext Ultra II DNA Library Prep Kit (NEB). Custom-made pyrrolo-cytosine adaptors (IDT) were substituted for kit-supplied adaptors. Adapted samples were bisulfite treated using the Premium Bisulfite Kit (Diagenode) and eluted in nuclease-free water. Post bisulfite clean-up, each sample was split into two 10 µL aliquots. One aliquot was further purified with a 1.2x left-side SPRI purification (Beckman). The SPRI-purified aliquot was denatured and APOBEC3A-treated (NEB, E1725L). After A3A deamination, the sample was purified via SPRI beads (1.2x, left-sided). Quantitative PCR was used to determine the number of cycles for the indexing PCR. Indexing PCR was performed using the unique dual index primer pairs from NEBNext Multiplex Oligos for Enzymatic-Methyl Seq (NEB). After the indexing PCR, samples were purified with two 0.8x left-sided SPRI purifications. Libraries were quantified with a Qubit HS kit (Invitrogen,Q32851) and assessed for quality with an Agilent Bioanalyzer. The resulting libraries were then pooled and sequenced on an Illumina MiSeq using 150-bp read length on a nano or micro sequencing kit (Illumina).

### EM-Seq library prep

Libraries were prepared using the NEBNext Enzymatic Methyl-seq kit (NEB, E7120L) in accordance with the manufacturer’s instructions. Either 200 ng or 20 ng of mouse gDNA with unmodified lambda and CpG-methylated pUC19 control DNA was sheared to 350 bp using a Covaris sonicator and used as input. HiFi HotStart Uracil+ Ready Mix (KAPA Biosystems, KK2801) was used to amplify the libraries following conversion and then subjected to a 0.8x left-sided SPRI purification, with elution in nuclease-free water to yield final libraries. Libraries were quantified by Qubit HS kit (Invitrogen, Q32851) and assessed for quality with an Agilent Bioanalyzer 2100. The resulting libraries were then sequenced on an Illumina MiSeq.

### Sequencing data processing

Sequencing data were processed in single-end mode as previously described [32]. Raw reads were trimmed for base quality and adaptor sequences using Trim Galore! (https://www.bioinformatics.babraham.ac.uk/projects/trim_galore/) and reads below a length of 20 bases were discarded. Pre- and post-trimmed data were visually examined for quality using FastQC (https://www.bioinformatics.babraham.ac.uk/projects/fastqc/). The trimmed reads were aligned to the reference genomes using Bismark (v0.22.3) with the following parameters -D 15 - R 2 -L 20 -N 0 –score_min L,0,-0.2 [42]. To account for A3A failure due to double-stranded DNA, reads with three or more consecutive non-converted cytosines in the CpH context were removed using the filter_non_conversion script included in the Bismark package. CpG sites from the sequencing data were classified based on chromatin states and aggregated. For each replicate at each timepoint, the 5hmCpG levels within each chromatin state were calculated from the bACE-seq methylation levels. Unmodified cytosine levels were calculated from the frequency of C-to-T changes at CpGs, while the 5mCpG levels were determined by subtracting the bACE-seq levels (5hmC) from the BS-seq modification levels (5mC and 5hmC together). For each replicate and each time point, the TAE was determined using the TAE calculator (https://zhou-lab.github.io/TAE_calculator/).

### Statistical analysis

A one-way ANOVA followed by Tukey’s range test was performed in R to compare methylation levels detected by different methods. Statistically significant differences in pair-wise comparisons are indicated by lowercase letters; groups with significant differences are assigned different letters, while groups with no statistical difference are assigned shared letters. n is defined as technical replicates. Exact values of n, significance, and effect size are reported in the text and figure legend.

## RESULTS

### Computational analysis of coverage-dependent precision in quantification of cytosine modifications

Although sparse sequencing of base-resolution libraries is commonly done to quality check libraries before high-depth sequencing, the reliability of the method for estimating 5mC and 5hmC levels and the relationships between coverage depth and accuracy are unknown. While sequencing continues to become more economical, in the absence of knowing the tradeoffs between depth and accuracy, it is not possible to construct rational strategies for using sparse sampling to study large cohorts of samples or to characterize rare cell types that might be inaccessible to methods demanding high sample input. We therefore began with a computational approach to assess the theoretical viability of Sparse-Seq pipelines before embarking on experimental endeavors. In this approach, to broadly profile across chemical and enzymatic methods and to be inclusive of both 5mC and 5hmC, we gathered data sets with whole-genome profiles available (genomic depth, range 1X to 9.5X) using various epigenetic pipelines (BS-Seq, TAB-Seq, ACE-Seq, and EM-Seq) in multiple murine and human cell types (**Figure 2, Figure S1, Figure S2**). We theorized that samples with lower overall levels of DNA modifications, such as those profiling 5hmC, might require greater coverage for accurate determination of cytosine modification levels, so we ensured that the data sets we chose had a range of overall modification levels, including mouse embryonic stem cells (ESCs), where 5hmC can be <0.3% of the total cytosine content.

**Figure 2.**
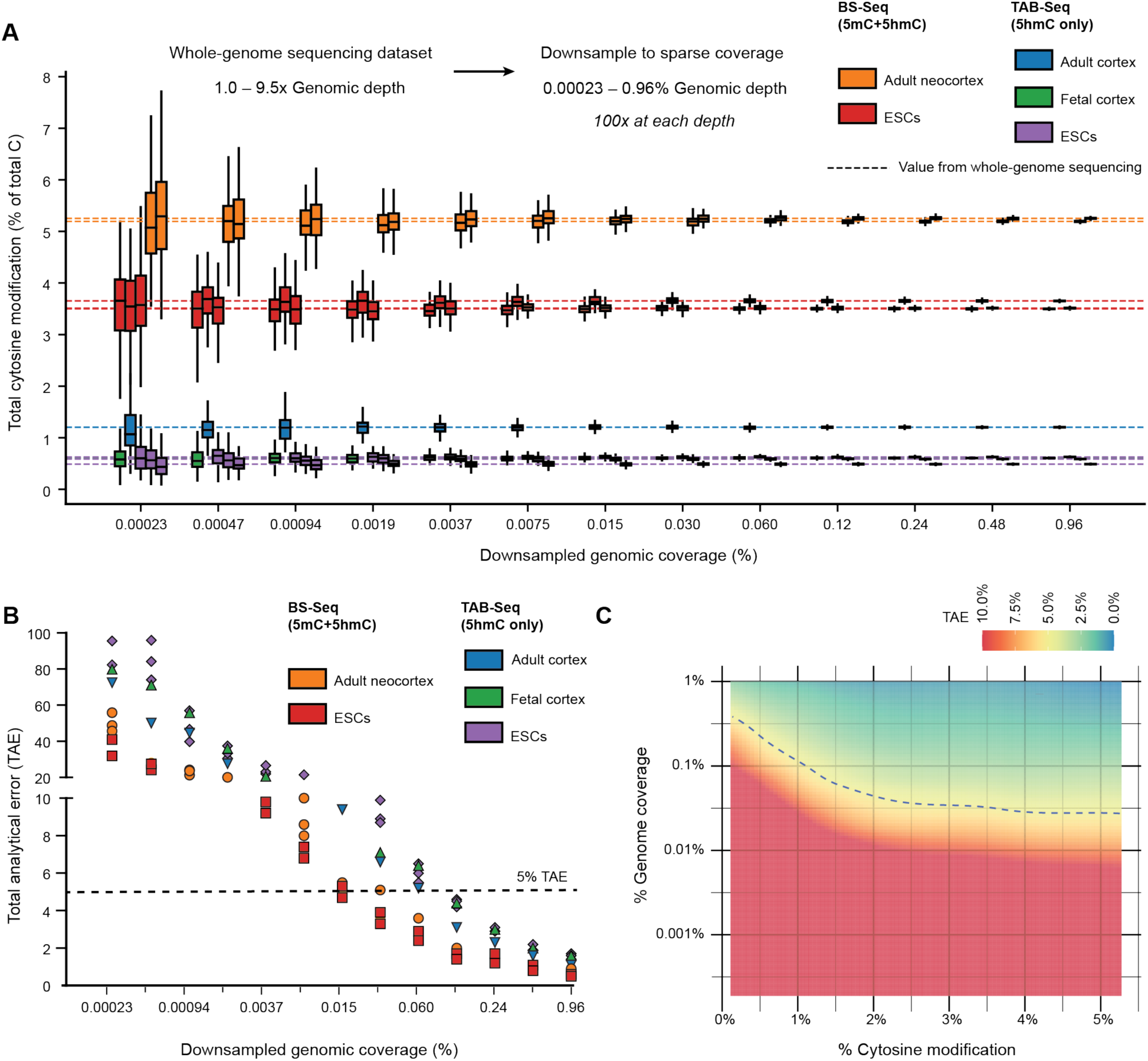
Computational downsampling confirms the theoretical viability of Sparse-Seq for cytosine modification measurement. **A)** Data from whole-genome sequencing in various murine cell types with either BS-Seq or TAB-Seq were randomly downsampled to various levels of sparse genomic coverage. Box plots represent the cytosine modification levels determined across 100 replicates at each sparse coverage level for each whole-genome dataset. Dashed lines mark the cytosine modification levels from the complete, deeply sequenced whole-genome dataset. Analogous analyses of whole-genome data sets obtained by enzymatic methods are shown in **Figures S1 and S2**. **B)** Shown is the calculated total analytical error (TAE) at each level of downsampling from the whole-genome datasets. The TAE gives the error in measurement of the true mean relative to the measured value for a single measurement at a given level of downsampling **C)** For 18 total whole-genome data sets (5 BS-Seq, 5 TAB-Seq, 4 ACE-Seq, and 4 EM-Seq), the genomic coverage in downsampling, the percent modification, and the associated TAE were taken as a base data set. LOESS regression was performed, and the regression model was then used to interpolate the TAE for genome coverage input ranging from 0.001% to 1% and total cytosine modification from 0% to 5%. The resulting model can be accessed in our TAE calculator tool at: https://zhou-lab.github.io/TAE_calculator/.

For each full genomic sequencing sample, we used the entire available high-depth data set to calculate the ‘true’ DNA modification level for 5mC+5hmC (BS-Seq, EM-Seq) or 5hmC alone (TAB-Seq, ACE-Seq). To model the process of our proposed Sparse-Seq, we then used computational downsampling to select a random subset of the complete sequencing reads. We targeted various values for genomic coverage, ranging from 0.00023% to 0.96% of the whole genome (e.g. ∼0.007 Mb to 26 Mb of the mouse genome). At each selected level of genomic coverage, downsampling was performed a total of 100 times, and these 100 sets of reads were separately processed to calculate the level of cytosine modification for each sample. We calculated the mean DNA modification level and standard deviation across the 100 sampled replicates at each level of genomic coverage. Because Sparse-Seq in practice would involve a single sampling at a given level of genomic coverage, we also calculated the total analytical error (TAE, % bias + 1.96 x coefficient of variance), a metric that assesses the expectation of test sample accuracy relative to the mean when run on a sample a single time.

The results show that across the epigenetic sequencing pipelines, even at the lowest genomic coverage we tested (0.00023%), the mean DNA modification level estimated across the 100 downsamplings was in reasonable agreement with the ‘true’ level from the complete dataset

(**Figure 2A**). As anticipated, however, variation is large at low levels of genomic coverage, but measurements converge as the level of genomic coverage increases (**Figure 2B**). For example, at 0.0019% genomic coverage, the BS-Seq and TAB-Seq datasets on both mouse ESCs and neocortex have an average TAE across samples of 24.8% (ranging from 14.2 to 37.4%). By contrast, when the genome coverage increases to 0.24%, the same datasets show a reduced TAE of 2.2% (ranging from 1.2 to 3.2%). This implies that if a single sample was profiled at 0.24% in sequencing depth, there would be a 95% probability that the determined level of cytosine modification will deviate by less than 2.2% from the ‘true’ value, as defined by high-coverage, whole-genome sequencing. By comparing across methods, we observed similar patterns across chemical and enzymatic-based sequencing methods (**Figure S1, Figure S2**), highlighting that the accuracy of downsampling is independent of the method of library construction. In the computational analysis, the TAE of each dataset correlates negatively with the overall modification level of the full dataset, supporting our hypothesis that Sparse-Seq would be less precise in samples with lower overall modification (at 0.24% coverage, TAB-Seq adult neocortex (5hmC = 1.2%) has a TAE of 2.3%, while TAB-Seq of ESCs (5hmC = 0.49%) has a TAE of 2.9%, **Figure 2B**). Nonetheless, even with lower overall levels of DNA modification, single sequencing replicates at 0.24% of the genome are associated with a TAE <5% for measurement of either 5mC+5hmC or 5hmC alone across all methods.

Our theoretical calculations with computational downsampling across different methods and samples thus isolated two key features, namely the percent coverage of the genome and the calculated mean DNA modification level, as instructive of the potential error associated with a single measurement of a given sample. Given the broad range of modification levels covered by our analysis (**Table S1**), we next used computational interpolation to construct a predictive landscape of TAE anticipated from measuring samples of different modification levels to various genome coverages (**Figure 2C**). As expected, the TAE prediction model integrating across the different data sets shows that the higher the modification level or genome coverage, the lower the TAE of the estimation. In anticipated applications of Sparse-Seq approaches, the end product of analysis would be a known percent coverage of the genome and an estimated modification level. Using the interpolation chart, the derived three-way relationship can be used to input genomic coverage and estimated modification level and output the expected TAE (**Table S2**). Alternatively, users can select a desired level of accuracy and use estimated modification levels to construct a target for the sequencing depth required to achieve that TAE.

To make error estimation with Sparse-Seq readily accessible, we developed an adaptable TAE calculator that allows users to determine the confidence of their Sparse-Seq measurements (https://zhou-lab.github.io/TAE_calculator/). Following low-coverage sequencing and alignment of samples, users simply provide three inputs into the calculator, the total read count, read length, and observed modification level (beta value) for their context of interest (total C, CpG, or CpH). The calculator then returns the expected TAE based on the interpolated model and reports the resulting beta value ± error (**Figure 2C**). This allows researchers to both prospectively design experiments to a desired accuracy threshold and retrospectively evaluate the reliability of completed measurements.

For many samples, the biological questions are centered on how levels of 5mC or 5hmC change in specific genomic elements (e.g., enhancers, promoters). In theory, given that reads are mapped in sparse sequencing approaches, they could also provide information on individual genomic elements, information that is inaccessible with LC-MS/MS based approaches. To explore this possibility, we first asked whether reads from whole-genome analysis are distributed proportionally to their prevalence in the genome, using the ChromHMM model for genomic annotation (**Figure S3**). We found that with the exception of some quiescent regions, likely enriched for repetitive elements, the distribution of reads in libraries accurately tracks the overall CpG prevalence of the reference genome. We thus postulated that the depth of sequencing needed to reliably report on genomic modification levels would increase for less prevalent modifications. Analogous to our genome-wide analysis, we plotted the TAE to estimate DNA modification levels for specific genomic elements. We found that sparse sampling could indeed reliably predict levels of 5mC and 5hmC in specific genomic elements including enhancers, gene bodies (e.g., transcribed regions), and repetitive elements or transcriptionally repressed chromatin (e.g., quiescent and polycomb repressed regions), with the relationship between TAE and depth following our predicted patterns (**Figure S4**). For quiescent elements, which make up the majority of the genome, the TAE curves nearly replicate those from whole-genome analysis. By contrast, for enhancers, which represent 0.44% of the genome, sampling to 8-fold greater depth was required to maintain similar TAE. Given that the TAE patterns for specific ChromHMM elements tracked with those of whole-genome data, we extended the TAE calculator to include analysis of specific genomic features. In this application of the calculator, the user can report on the modification level of interest (beta value) within a given genomic element. The calculator scales down the calculated genome wide coverage by the prevalence of the ChromHMM state in the genome and reports the associated TAE of the measured value.

Considering the generalizability of the above sparse sequencing approach in estimating cytosine modification levels and quantitatively controlling errors, we returned to the question of genome-wide levels and selected a target range of genomic depth coverages that would allow for accurate quantification and higher throughput analysis. A coverage level of 0.24% of the mouse genome is equivalent to sequencing approximately 6.5 million bases. With NGS read length of 150 bases, this level of genomic coverage would be equivalent to about 45,000 (45K) mapped reads. If users are focused on specific genomic elements, the depth of sequencing can be increased in a rational manner using the TAE estimator for a predicted level of accuracy. Given the wide range of flow cells available and the relative accessibility of sequencing to most laboratories, we considered that Sparse-Seq at these levels of coverage could open opportunities for economical and efficient sequencing for a wide variety of applications, particularly if the methodology showed comparable experimental performance to LC-MS/MS-based approaches.

### Comparison of Sparse-Seq with multiple sequencing pipelines to LC-MS/MS

Based on the results from our computational work, we next moved to experimentally explore Sparse-Seq for resolution of genome-wide cytosine modification levels using a target of ∼45K mapped reads per sample, with a rigorous comparison to LC-MS/MS based methods. Given that our objective was accurate quantification of both 5mC and 5hmC levels from genomic DNA samples, we chose to employ Sparse-Seq using the BS and bACE-Seq pipeline [30, 31]. This approach combines bisulfite and enzymatic deamination to parse 5mC and 5hmC (**Figure 3A)**. Specifically, genomic DNA is subjected to chemical deamination with bisulfite, which leads to the conversion of C, but not 5mC or 5hmC, separating unmodified C from modified C bases. Half of the sample is then subjected to enzymatic deamination with the DNA deaminase, APOBEC3A (A3A). A3A can readily deaminate the 5mC bases, but is unable to deaminate the 5hmC bases, which were converted to the bulky and deaminase-resistant base, cytosine 5-methylenesulfonate (CMS), after the bisulfite step. By comparison of the BS-Seq (5mC+5hmC read as C) with the bACE-Seq library (only 5hmC reads as C), the global level of both 5mC and 5hmC can be calculated. We focused on this approach as an accessible option for labs without access to LC-MS/MS given that BS-Seq is the most commonly employed approach for sequencing and the commercial availability of A3A.

**Figure 3.**
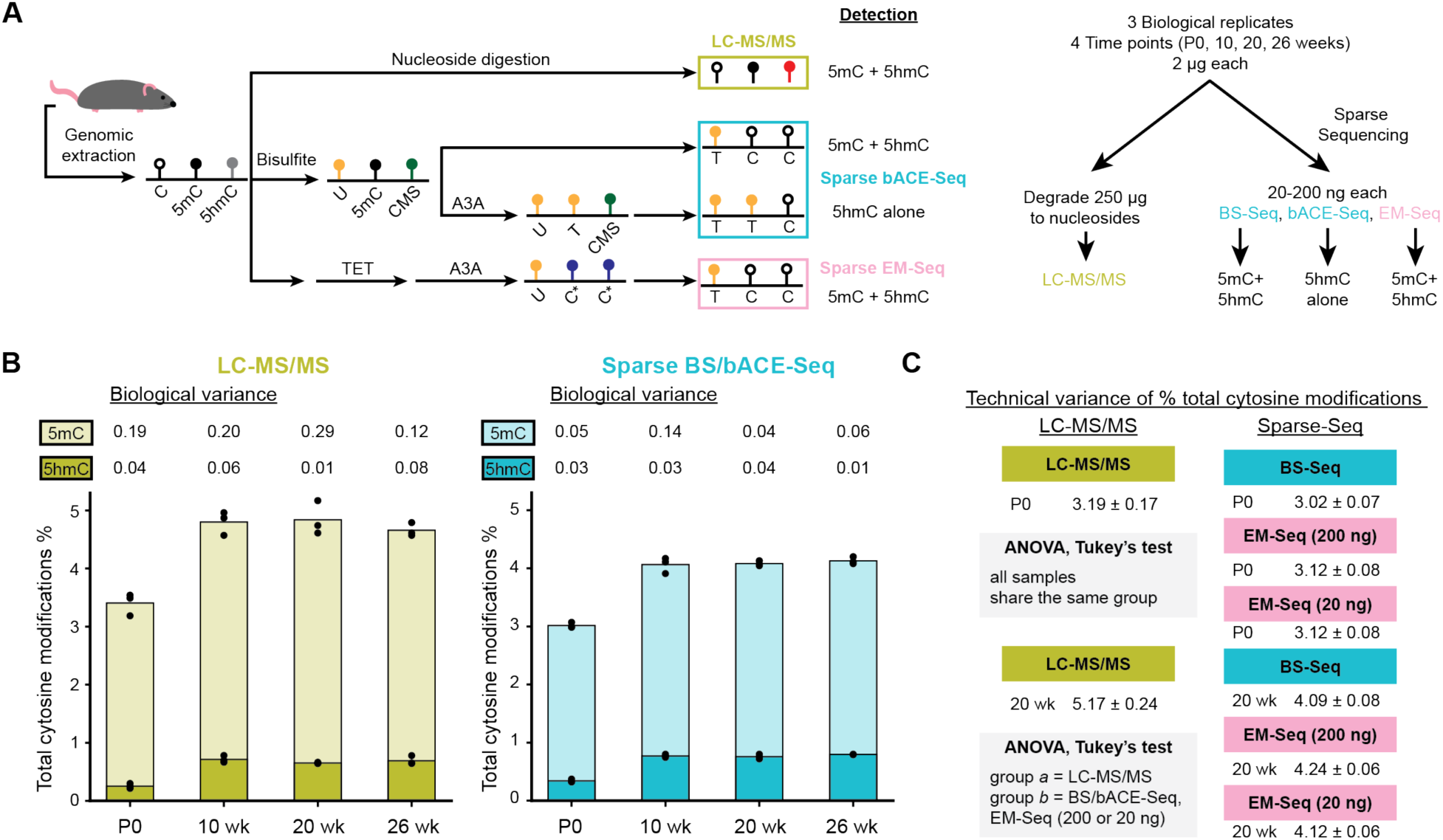
The accuracy, precision, and generalizability of Sparse-Seq is confirmed with comparison to LC-MS/MS. **A)** Workflow for experimental validation of Sparse-Seq using BS-Seq, bACE-Seq, and EM-Seq pipelines via comparison to LC-MS/MS using murine cortex gDNA. Abbreviations: CMS, cytosine 5-methylenesulfonate; C*, modified cytosine protected from enzymatic deamination. **B)** Biological variance comparison of 5mC and 5hmC quantification using LC-MS/MS (left) and BS/bACE-Seq (right). Each of the three biological replicates is shown (points), with bars representing the average of measurements and standard deviations listed above the bars. **C)** Technical variance comparison of LC-MS/MS and Sparse-Seq. Data are from three technical replicates from a single sample from (**B**) at either P0 and 20 weeks using LC-MS/MS, BS-Seq, or EM-Seq at either of two sample inputs (200 ng or 20 ng), with mean and standard deviation provided for each set of technical replicates. A one-way ANOVA with Tukey’s range test, n=3; significant differences between groups are denoted using different letters, while groups with no significant differences share the same letter.

To permit an assessment of biological and technical variation, we collected three independent biological samples of murine brain genomic DNA (gDNA) at four distinct time points in mouse development ranging from neonates (P0) to 26 weeks old (**Figure 3A**). We processed 250 ng of the gDNA with a standard LC-MS/MS protocol involving degradation to nucleosides and 200 ng of the gDNA with the Sparse BS and bACE-Seq pipeline, pooling the barcoded samples for analysis via NGS. For LC-MS/MS, the degraded gDNA was analyzed for detected masses and quantification was performed against a standard curve generated for each DNA modification. For NGS, we aimed for equivalent library pooling targeting similar depth across samples. We obtained a range of ∼71-151K reads (average 108K), which, after alignment and de-duplication, yielded ∼27-102K reads across samples (average 68K). Notably, while some samples thus fell below our goal of 45K, the TAE calculator predicted acceptable reliability (< 2.5% TAE). In comparing the global cytosine modification levels for both 5mC and 5hmC, we found strong concordance between the values determined using Sparse-Seq and LC-MS/MS (**Figure 3B**). Overall, the relative abundance of the individual modifications followed the same rank order for both methods, although Sparse-Seq appeared to consistently detect more 5hmC and less 5mC relative to the LC-MS/MS. Importantly, it is not clear which method would be more accurate in this regard. In LC-MS/MS, each modification is quantified against its own standard, which can lead to potential compounded errors from multiple standard curves. In Sparse-Seq, the modification levels are derived from the comparison of C versus T calls at each site, with errors that could instead be associated with incomplete conversions. Importantly, across samples, LC-MS/MS showed consistently higher variance in 5mC quantification than Sparse-Seq across the same samples, by a range of 1.5 to 6-fold. This difference is likely attributable to our ability to precisely control TAE in Sparse-Seq by targeting a sequencing depth that met an acceptable TAE threshold for our application. To assess for technical variation, we repeated LC-MS/MS analysis and Sparse-Seq on a set of the biological replicates and observed higher consistency with Sparse-Seq, with a variance that was 2.1- to 4.0-fold lower than LC-MS/MS in 5mC detection (**Figure 3C**). The higher degree of technical variance observed in measurements from LC-MS/MS suggests at least one source of the greater overall variance observed with biological replicates using LC-MS/MS relative to Sparse-Seq.

Given the increased momentum towards enzymatic rather than chemical approaches to detect cytosine modifications [10], we additionally explored Sparse-Seq with an alternative all-enzymatic sequencing approach. We chose EM-Seq due to its commercial accessibility as a method for quantifying combined 5mC+5hmC levels, also recognizing that this orthogonal approach could help assess the relative accuracy of BS-Seq versus LC-MS/MS given their slight discordance. An added benefit of enzymatic library generation approaches, such as EM-Seq and ACE-Seq, is that these approaches require much lower starting input amounts of gDNA than either LC-MS/MS methods or bisulfite-based methods. To this end, given our interest in the technical variance of each method, we processed one biological gDNA sample from P0 and another from the 20-week timepoint with LC-MS/MS (250 ng input gDNA), Sparse BS-Seq (200 ng), and Sparse EM-Seq using either 200 ng or 20 ng input gDNA. After processing, the BS-Seq and EM-Seq data sets had a range of 10-90K reads (average 56K) which were used to calculate the levels of 5mC + 5hmC. By performing Tukey’s HSD Test for multiple comparisons, we found that the two orthogonal sequencing methods (Sparse BS-Seq vs. EM-Seq) agreed more closely with each other than LC-MS/MS (p < 0.0001) with a large effect size (η^2^=0.94) (**Figure 3C**). While all methods will have some biases, the consistency between the different enzymatic and chemical-based sequencing methods, relative to LC-MS/MS, highlights possible advantages in the accurate detection of modification levels. Regarding technical variance, LC-MS/MS showed 4.6-5.3% variability between independent technical measurements, while 1.9-2.5% variability was observed across Sparse-Seq samples. Overall, across analysis of biological and technical reproducibility, the data highlight the generalizability of Sparse-Seq across chemical and enzymatic methods as well as its higher precision relative to LC-MS/MS at the tested sequencing depths.

### Sparse-Seq reveals distinct developmental timing of cytosine modifications

One of the most notable disadvantages of LC-MS/MS for profiling cytosine modifications is that the approach erases all information on sequence context given the requirement for degradation of gDNA to nucleosides. While Sparse-Seq lacks whole-genome base-resolution information, it does allow for analyses of sequence context (e.g. CpG versus CpH) or mapping of DNA modification levels to specific genomic elements.

To explore the utility of Sparse-Seq in addressing open biological questions, we focused analysis on the developing brain. Prior work has shown that during development, two different DNA marks accumulate on cytosine bases in post-mitotic neurons. First, there is an accumulation of 5mC in non-CpG (CpH) contexts [43], and second, an accumulation of 5hmC, confined almost exclusively to CpG contexts [43, 44]. Notably, based on single-cell profiling work, either 5mCpH alone or 5hmCpG alone can be used to define cell identity and lineage in the brain, highlighting the importance of both of these epigenetic marks [31, 45]. While both accumulate during brain development, the challenge and expense of base-resolution mapping have meant that the relative kinetics with which 5mCpH and 5hmCpG arise during prenatal and postnatal brain development have been incompletely characterized. Furthermore, while 5hmC is known to accumulate, the relative kinetics of changes in 5mC versus 5hmC in different genomic elements is not known.

Given that Sparse-Seq is well-situated to assess DNA modification levels while retaining information on sequence context and genomic elements, we collected murine brain gDNA samples at various points in development from E16 to 10 weeks of age, with 3 biological replicates for each timepoint (**Figure 4A**). We then subjected the samples to the split Sparse BS-Seq and Sparse bACE-Seq approach to quantify both 5mC and 5hmC. With a primary goal of mapping global modification levels while balancing cost and accuracy, we targeted ∼30K mapped reads and ran the samples on an Illumina Micro kit (resulting range: 11–54K mapped reads, average 29K).

**Figure 4.**
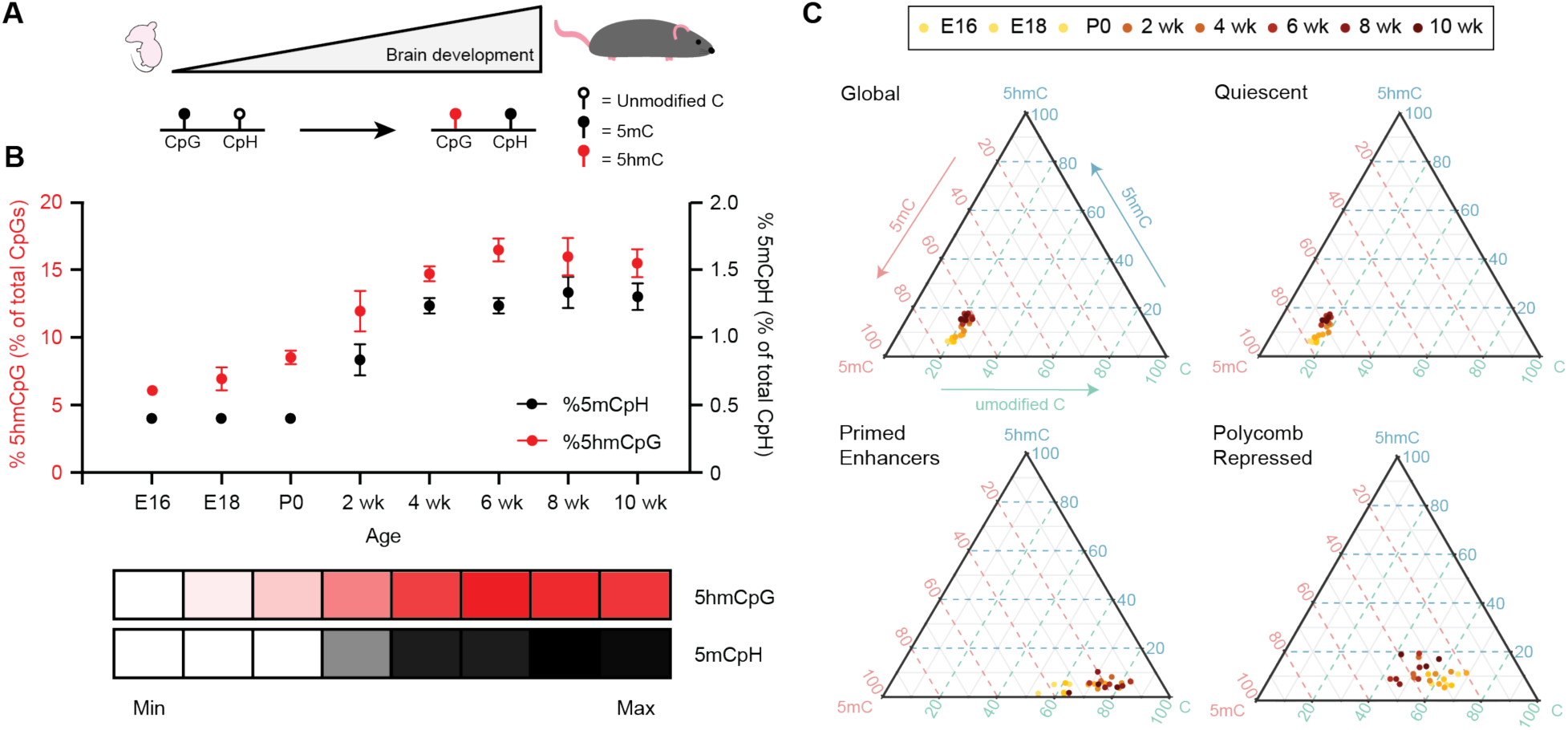
Sparse-Seq reveals 5mCpH and 5hmCpG dynamics during murine development. **A)** The developing mouse brain is known to accumulate 5mC in non-CpG (CpH) contexts and 5hmC in CpG contexts. Three biological replicates of brain specimens (whole brain for fetal and P0 stages and isolated cortex thereafter) taken every two weeks from E16 through 10 weeks were analyzed by Sparse BS/bACE-Seq and quantified for the level of modifications in each context. **B)** Levels (%) of 5mCpH and 5hmCpG in the developing mouse brain detected by Sparse BS/bACE-Seq. Data from the replicates at each time point are shown as the mean, with standard deviation provided by error bars. Error bars are not shown if they are smaller than the symbol representing the mean value. At the bottom is a heat map depicting the relationship between 5hmCpG and 5mCpG from minimal to maximal values over time. **C)** Shown are ternary plots representing the level of C, 5mC, and 5hmC measured for the whole genome or specific genomic elements. Each data point represents the measurement from a single biological replicate, with different time points represented by different colors.

Consistent with prior studies, an accumulation of both methylation in CpH contexts (5mCpH) and hydroxymethylation in CpG contexts (5hmCpG) was observed. Tracking these modifications at more regular intervals between embryonic development and young adulthood with our Sparse-Seq approach yielded several new insights. As anticipated, both 5mCpH and 5hmCpG accumulate with development. However, the kinetic patterns of accumulation can differ. First, a surprisingly high amount of 5hmC can be observed in CpG contexts even in the earliest developmental time point collected, E16 (**Figure 4B**), consistent with an early role for TET enzymes in shaping brain epigenetic landscapes [7, 13, 14]. By birth (P0), 25% of the total 5hmC accumulation observed between E16 and 10 weeks of age has already occurred, after which levels continue to rise. By contrast, non-CpG methylation (5mCpH) is at background levels at birth (P0), with a rapid increase that occurs post-partum. Interestingly, both 5hmCpG and 5mCpH levels, however, appear to reach 90% of their maximal levels by 4 weeks after birth, suggesting that the epigenetic marks are largely established at that point in brain development. The shared and divergent patterns with which these epigenetic marks arise highlight the likelihood of independent epigenetic roles for 5hmC and non-CpG methylation in neuronal development. Notably, these insights are supported by taking the mean and variation from merged biological replicates (**Figure 4B**), but the strength of conclusions can also be ascertained by assessing the variability of individual biological observations, each with their own associated TAE (**Figure S5**), demonstrating the value of using a rationally determined target sequencing depth for the study of a large cohort of samples.

We next turned our attention to tracking dynamic changes in 5mCpG and 5hmCpG at specific genomic elements. To observe the changes with time, we plotted the quantified levels of C, 5mC, and 5hmC in each genomic element in a ternary plot, where each data point includes the level of all three modification states, which sum to unity. These plots could readily convey that the DNA methylation dynamics can differ greatly by genomic element (**Figure 4C, Figure S6**). For example, the levels of DNA modifications in quiescent regions, which make up the majority of the genome, track with global changes. Unmodified C levels are generally constant across time from E16 to week 8 (16.2 ± 1.3% to 16.4 ± 0.3%), while 5hmC accumulates (6.0 ± 0.3% to 15.8 ± 1.3%) and a concomitant decrease in 5mC (77.8% ± 1.0% to 68.7 ± 1.4%). By contrast, at select enhancers, we instead observe that 5hmC levels are low and largely unchanged. At these elements, unmodified C accumulates during the prenatal period (57.6 ± 4.5% to 75.4 ± 5.8%), with the change accounted by the loss of 5mC over the same time (38.5 ± 6.4% to 18.3 ± 3.5%). A reverse of this pattern can be observed at polycomb repressed regions, where there is a major gain in 5mC (28.9 ± 5.7% to 42.3 ± 6.1%), while unmodified C decreases (63.1 ± 4.0% to 47.9 ± 3.0%), concurrent with some increase in 5hmC. Notably, as our target sequencing depth was selected with the initial goal of studying global 5hmCpG and 5mCpH levels, the measurements of modification levels within individual ChromHMM elements would be expected to have higher TAE. To better characterize these effects, we focused on the ChromHMM element associated with active transcription (Tx). For each observation at a given time point, we calculated the levels of unmodified CpG, 5mCpG, and 5hmCpG and used our TAE calculator to derive the associated error. As Tx represents ∼4.7% of the overall genome, the associated TAE values were indeed significantly higher (range 7.6% to 35%). Nonetheless, despite the higher TAE with individual data points, we could definitively observe that 5hmCpG increases in Tx in the post-natal setting, with concomitant decrease in 5mCpG and minimal change in unmodified CpGs (**Figure S7**). Thus, even from Sparse-Seq data with a read-depth targeted to accurately track global DNA modification levels, the trends in changes in genomic elements can be readily ascertained.

## DISCUSSION

The ability to accurately, efficiently, and economically estimate the level of DNA modifications in samples derived from cells or tissues is of high value. The levels of DNA modifications can be used to track cellular and developmental dynamics or to classify, diagnose, or prognosticate disease states. While multiple methods exist for obtaining estimates of 5mC and 5hmC levels, the accessibility of sequencing makes Sparse-Seq a potentially generalizable solution to address these needs. Our goal was to computationally establish a strong theoretical foundation for low-coverage sequencing, develop user-friendly tools for implementation of Sparse-Seq, experimentally validate its performance against LC-MS/MS, and apply the method to derive new biological insights that benefit from preserving sequence context in analytical pipelines.

While sequencing is a cornerstone of genomics, LC-MS/MS-based approaches are currently generally regarded as the ‘gold standard’ for estimation of global levels of 5mC and 5hmC. Here, to directly compare these methods and evaluate Sparse-Seq, we performed comparisons to LC-MS/MS, with our results indicating comparable quantification of C, 5mC, and 5hmC with less technical variability. Although our results were generally concordant, we noted systematic differences in certain measurements. We postulate that with LC-MS/MS, systematic errors might be amplified. A potential explanation for LC-MS/MS detecting less 5hmC and more 5mC than Sparse-Seq is that each modification is quantified against its own separate standard curve, which may introduce compounding errors across modifications. It is feasible that the accuracy of LC-MS/MS could have been further increased in our study by the use of internal isotopic controls for each nucleoside, rather than external calibration controls [46]; however, this approach is inconsistently used in the literature, given added cost and more limited accessibility to modified nucleoside isotopologues. In sequencing-based approaches, modification levels are derived from internal comparisons, specifically the number of bases that read as C versus T at each aligned base across reads. If deeply sequenced whole-genome data sets are regarded as the most reliable means to estimate global modification levels, integrating across our computational and experimental data, a case can thus be made for Sparse-Seq as a viable alternative to LC-MS/MS methods.

The derivation of mappable reads is a distinctive advantage of Sparse-Seq approaches over LC-MS/MS, as it preserves sequence context and allows for segregation by genomic elements. In applying Sparse-Seq to dissect cytosine modification dynamics in the developing mouse brain, we were able to analyze the time course with which 5hmCpG and non-CpG methylation occur. While both modifications accumulate in a time-dependent manner and plateau before 6 weeks after birth, we find that 5hmC starts to accumulate in the prenatal brain, while non-CpG methylation accumulation only begins postnatally. These non-CpG methylation changes could be explained in part by the temporal pattern of DNMT expression, with evidence suggesting that DNMT3A is specifically induced in the post-natal period, peaking at 3 weeks before declining [47]. Additionally, when the data were segregated by genomic element, we were able to track distinct patterns of cytosine modification dynamics, with compelling examples where trade-offs can be entirely inversed. For example, while 5hmC levels accumulate globally, they are generally low and stable in both polycomb repressed regions and primed enhancers. At these genomic elements, a gain of methylation and loss of unmodified C can be observed in the polycomb repressed regions, while the opposite pattern is evident in primed enhancers. Notably, if the goal of a Sparse-Seq experiment were to focus on specific genomic elements, one can adjust the depth of sequencing to allow for any given level of acceptable error by using the TAE calculator. On a related note, an added strength of Sparse-Seq is that the library preparation is identical to that needed for high-depth sequencing. Thus, samples identified as being of high value after assessment of global modification levels, for example, the timepoints at which 5mC and 5hmC are most rapidly changing, can be preserved and sequenced at high depth for base-resolution analysis.

Sparse-Seq offers a flexible method that can be used with different sequencing workflows to quantify desired modifications of interest with tunable sensitivity. We demonstrate that the method can be applied to resolve combined levels of 5mC + 5hmC (e.g. BS-Seq, EM-Seq), or integrated to yield 5mC and 5hmC alone (e.g. TAB-Seq, bACE-Seq). Importantly, the extension to all-enzymatic approaches could be particularly valuable for the study of samples where DNA amounts are limiting. Enzymatic methods, such as EM-Seq and DM-Seq, are typically performed with low quantities of DNA at the nanogram level and have even been extended to the single-cell level, offering opportunities not readily accessible to LC-MS/MS [33, 35, 48]. One associated limitation of Sparse-Seq is that the detection limit for given modifications can be impacted by both the accuracy of correct base calling in NGS as well as the rate of failures in proper C-to-T conversion for a given sequencing pipeline. However, these rates can typically be accurately estimated, for example, by examining the frequency of BS-Seq failure in the CpC context, where methylation is not commonly found in mammalian genomes. Additionally, as the frequency of a given modification determines the depth of sequencing needed, our data suggest that while Sparse-Seq is now validated for studying unmodified C, 5mC and 5hmC, it is not as well suited for quantification of exceptionally rare DNA modifications such as 5-formylcytosine or 5-carboxycytosine, whose levels can be 1 in 10^5^ of all cytosines or even less [4, 7]. Notably, recent work has also suggested the possibility of sparse sampling of DNA methylomes through nanopore approaches, which may have different associated accuracy errors [49, 50]. Although the validity of parsing 5hmC by nanopore has yet to be rigorously established, some of the principles we explored in short-read sequencing platforms could likely be extended to long-read approaches with sparse sampling.

With a theoretical foundation, experimental validation, and an accessible TAE calculator now established, it is important to consider how Sparse-Seq should be situated within the landscape of existing sequencing technologies (**Figure S8**). Although many scenarios can be imagined, we anticipate that the method will be most valuable for biologists at early stages of investigation or studying larger cohorts of samples. Here, key questions often precede the commitment to expensive deep sequencing, such as: “Are cytosine modification levels in fact changing?”; “Is this occurring in a subset of the samples or at specific time points?”; and “Can such changes be localized to specific genomic features?”. Sparse-Seq offers an economical means to survey across samples, either to selectively target sequencing depths for firm conclusions to be drawn or to rationally nominate a subset of samples for follow-up high-depth sequencing studies.

While we profiled serial time points during development in our example, the approach is more broadly applicable. Compelling use cases include characterizing large cohorts of samples for changes in 5mC or 5hmC as a function of tumor stage, profiling 5mC and 5hmC levels across multiple tissues or cell types from diverse species, or deciphering how various environmental stressors can alter 5mC or 5hmC landscapes in different genomic elements. Given the ubiquity of sequencing relative to LC-MS/MS, the compatibility of Sparse-Seq with a broad array of chemical and enzymatic methods, and the availability of a TAE calculator, we propose that Sparse-Seq should now be a first-line consideration whenever global or element-specific quantification of 5mC and 5hmC is the initial goal.

## DATA AVAILABILITY

Analysis performed using previously developed software is detailed in the relevant methods section. The TAE calculator is publicly available (https://zhou-lab.github.io/TAE_calculator/). All original code used for analyses in this paper have been deposited on GitHub (https://github.com/ChristianELoo/Sparse-Seq) and are publicly available as of the date of publication. NGS data have been deposited at GEO (GSE273521) and are publicly available as of the date of publication. For reviewers, GEO accession GSE273521 can be accessed via: https://www.ncbi.nlm.nih.gov/geo/query/acc.cgi?acc=GSE273521, with token edebiomsxhqtbwt. Additional information required to reanalyze the data reported in this paper is available from the lead contact upon request.

## SUPPLEMENTARY DATA

Supplementary Data are available at NAR online.

## AUTHOR CONTRIBUTIONS

Conceptualization, C.E.L, J.M.F., W.Z., and R.M.K.; Methodology, C.E.L., J.M.F., H.Z., W.Z., R.M.K.; Software, W.Z.; Investigation, C.E.L., J.M.F., H.Z., R.Z.; Writing - Original Draft, C.E.L., J.M.F., R.M.K.; Writing - Review and Editing, All Authors; Resources, M.S.B., C.K., W.Z., R.M.K.; Supervision – W.Z. and R.M.K.

## ACKNOWLEDGEMENTS

We thank Tong Wang for guidance and Robert J. Ontiveros for providing LC-MS/MS training.

## SUPPLEMENTARY INFORMATION

### Supplementary Figures

**Figure S1.**
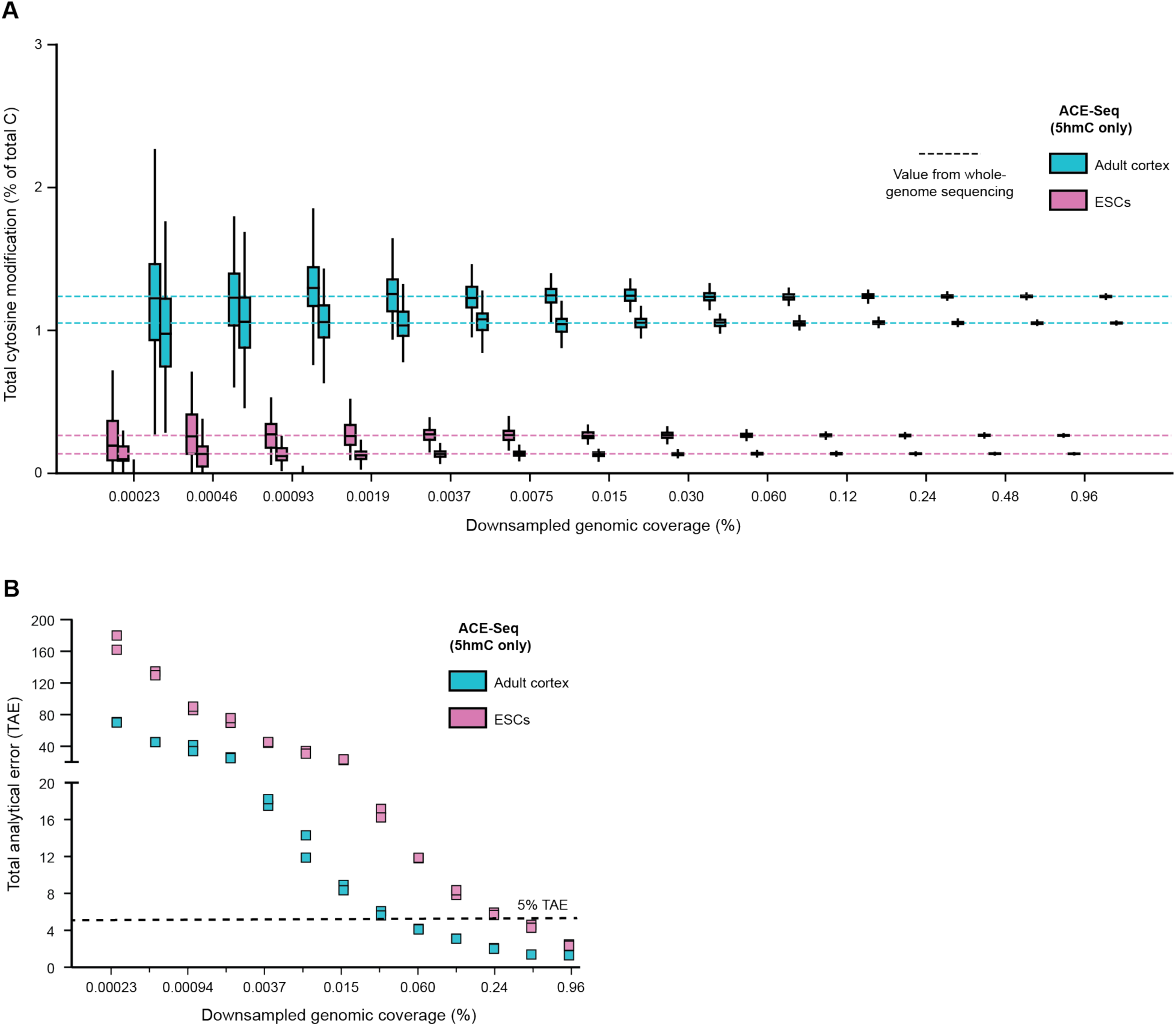
Computational downsampling applied to published ACE-Seq datasets. **A)** Whole-genome sequencing data sets sequenced with ACE-Seq in various murine cell types were downsampled to sparse genomic coverage. Box plots represent the cytosine modification levels determined in each of 100 replicates at each sparse coverage level. Dashed lines mark the cytosine modification levels from the complete, deeply sequenced whole-genome dataset. **B)** Shown is the calculated total analytical error (TAE) as a function of the downsampled coverage from the whole-genome datasets.

**Figure S2.**
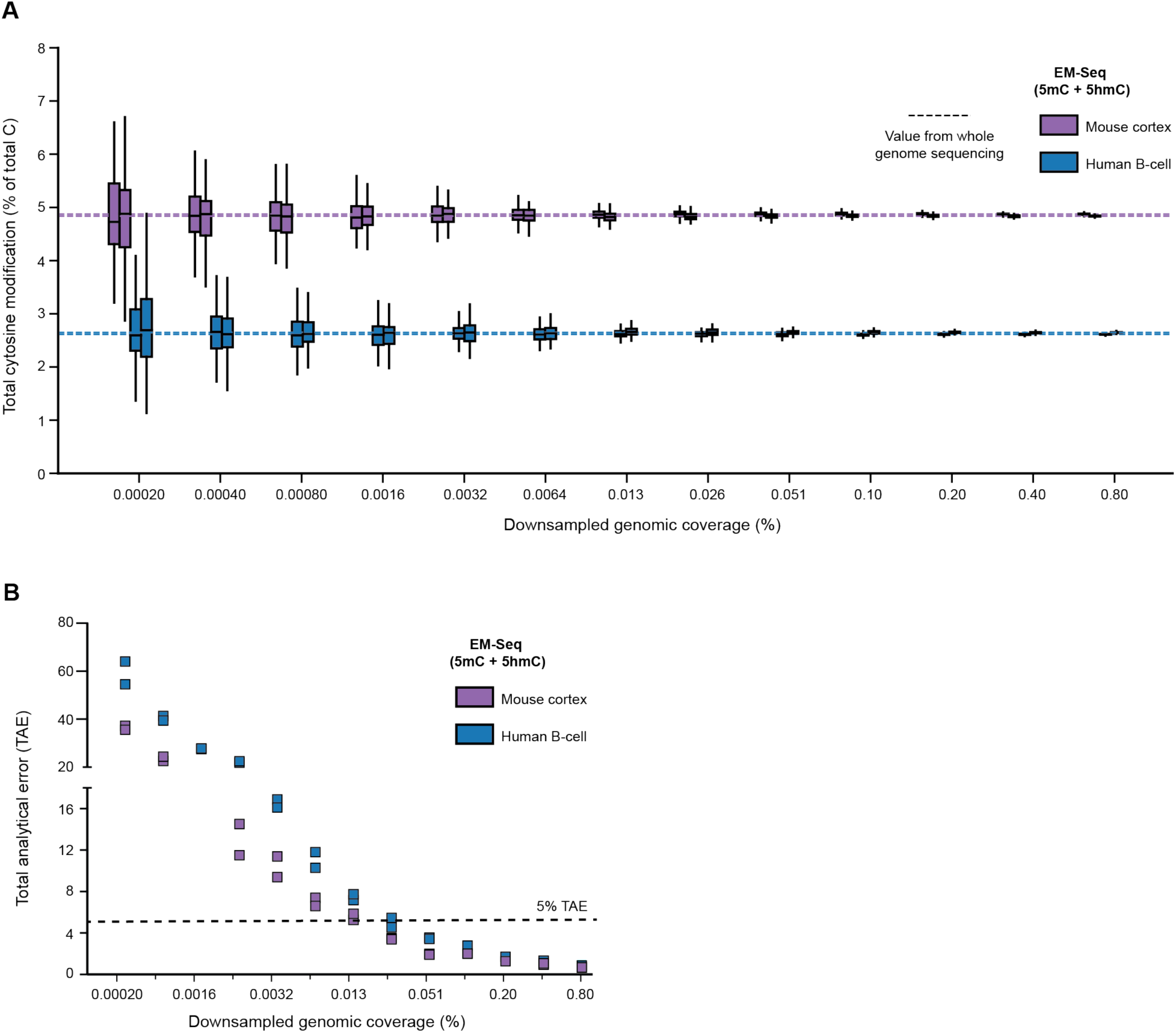
Computational downsampling applied to published EM-Seq datasets. **A)** Whole-genome sequencing data sets sequenced with EM-Seq in various murine cell types were downsampled to sparse genomic coverage. Box plots represent the cytosine modification levels determined in each of 100 replicates at each sparse coverage level. Dashed lines mark the cytosine modification levels from the full, deeply sequenced whole-genome dataset. **B)** Shown is the calculated total analytical error (TAE) as a function of the downsampled coverage from the whole-genome datasets.

**Figure S3.**
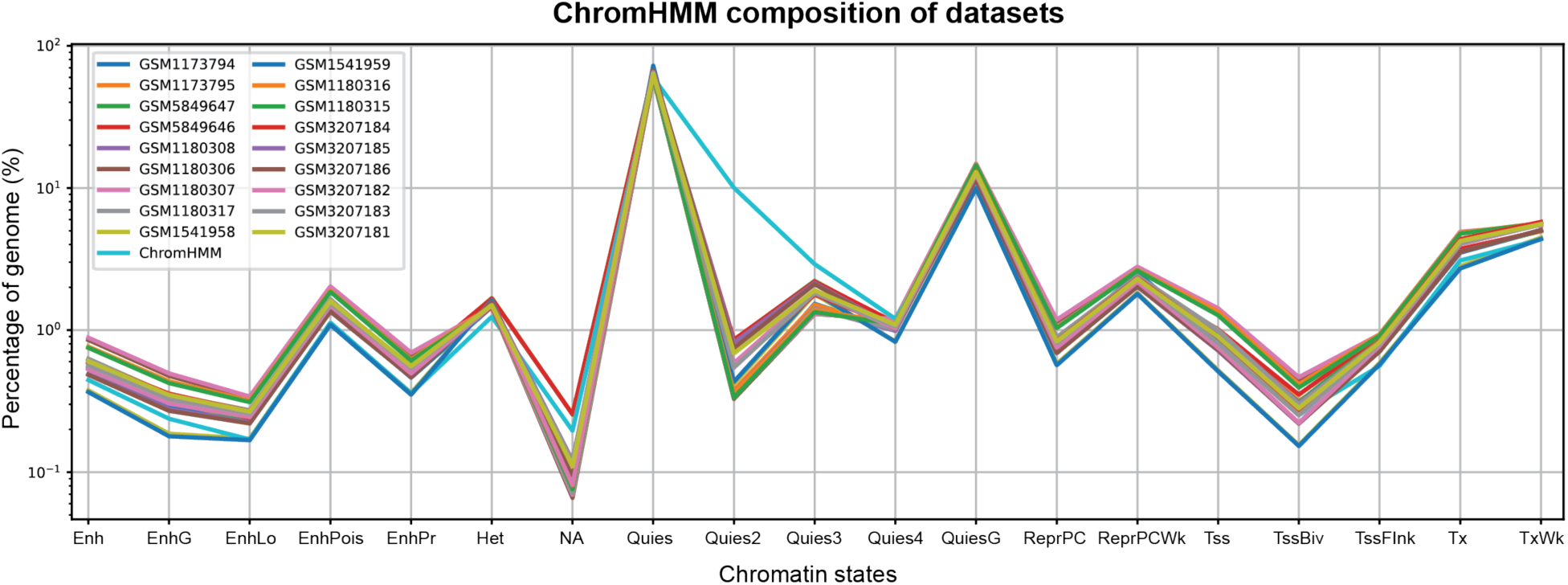
Distribution of genomic element prevalence relative to mappable reads. Using various genomic elements categorized by ChromHMM, plotted in light blue is the percentage of the genome (log-scale) attributable to a given genomic element. For the other deeply sequenced, whole-genome datasets, plotted are the percentage of reads that uniquely map to a given genomic element. While fewer reads map to quiescent regions, for other genomic elements the percentage of reads mapped from deeply sequenced data sets generally tracks with their abundance in the genome.

**Figure S4.**
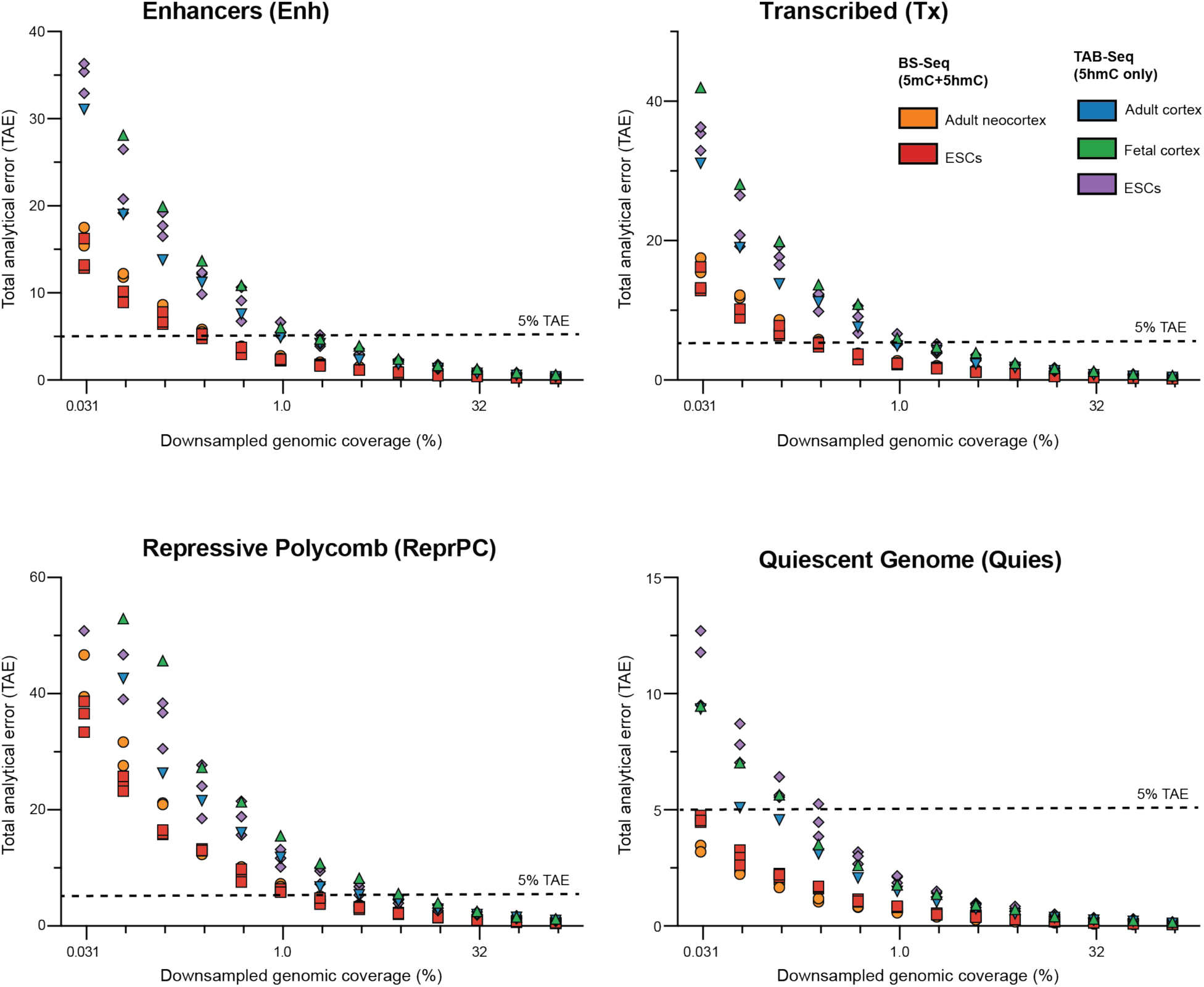
Computational downsampling applied to genomic element analysis for whole-genome datasets. Whole-genome sequencing datasets generated from various methods were downsampled to sparse genomic coverage. The resulting reads were mapped and categorized by ChromHMM element, correlating with different common genomic elements. Shown is the calculated total analytical error (TAE) as a function of the downsampled coverage for various genomic elements.

**Figure S5.**
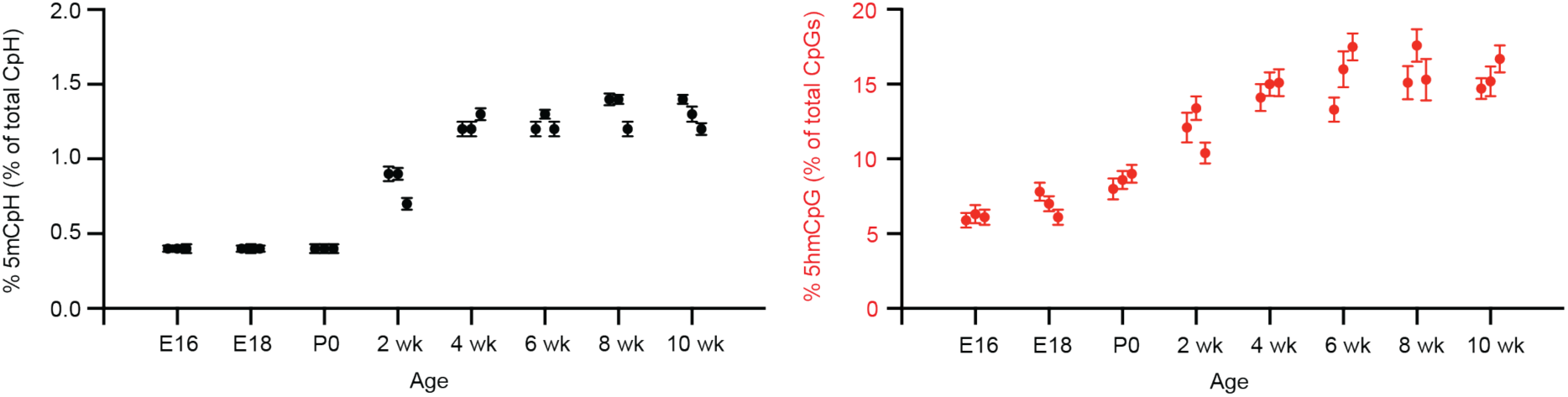
Total Analytical Error (TAE) for 5mCpH and 5hmCpG measurements across murine development. Shown are plots of 5mCpH (left) and 5hmCpG (right) levels (%) in the developing mouse brain detected by Sparse BS/bACE-Seq. At each timepoint, three data points are shown, each representing an independent biological replicate. TAE bars derived from the TAE calculator are shown for each datapoint.

**Figure S6.**
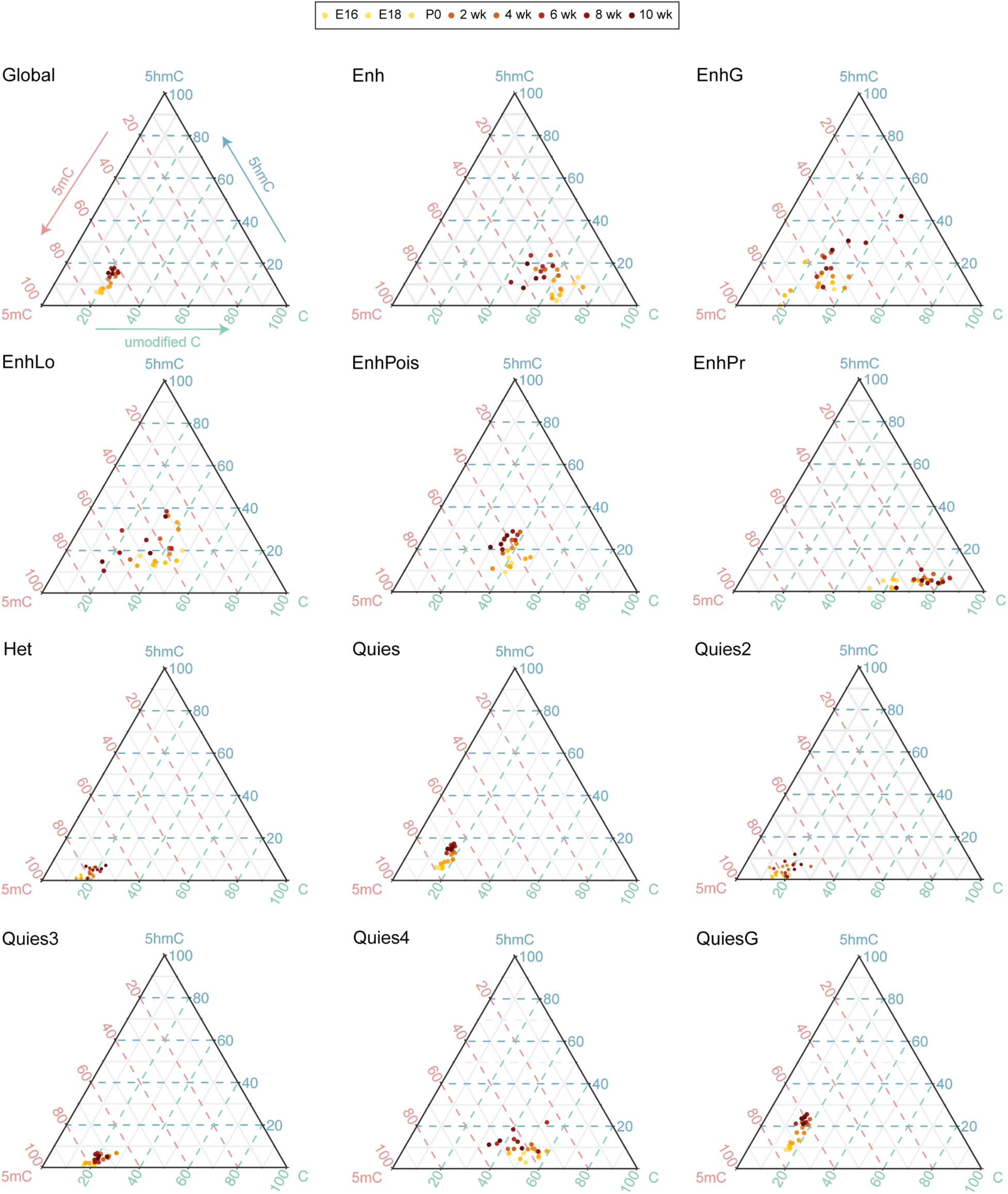
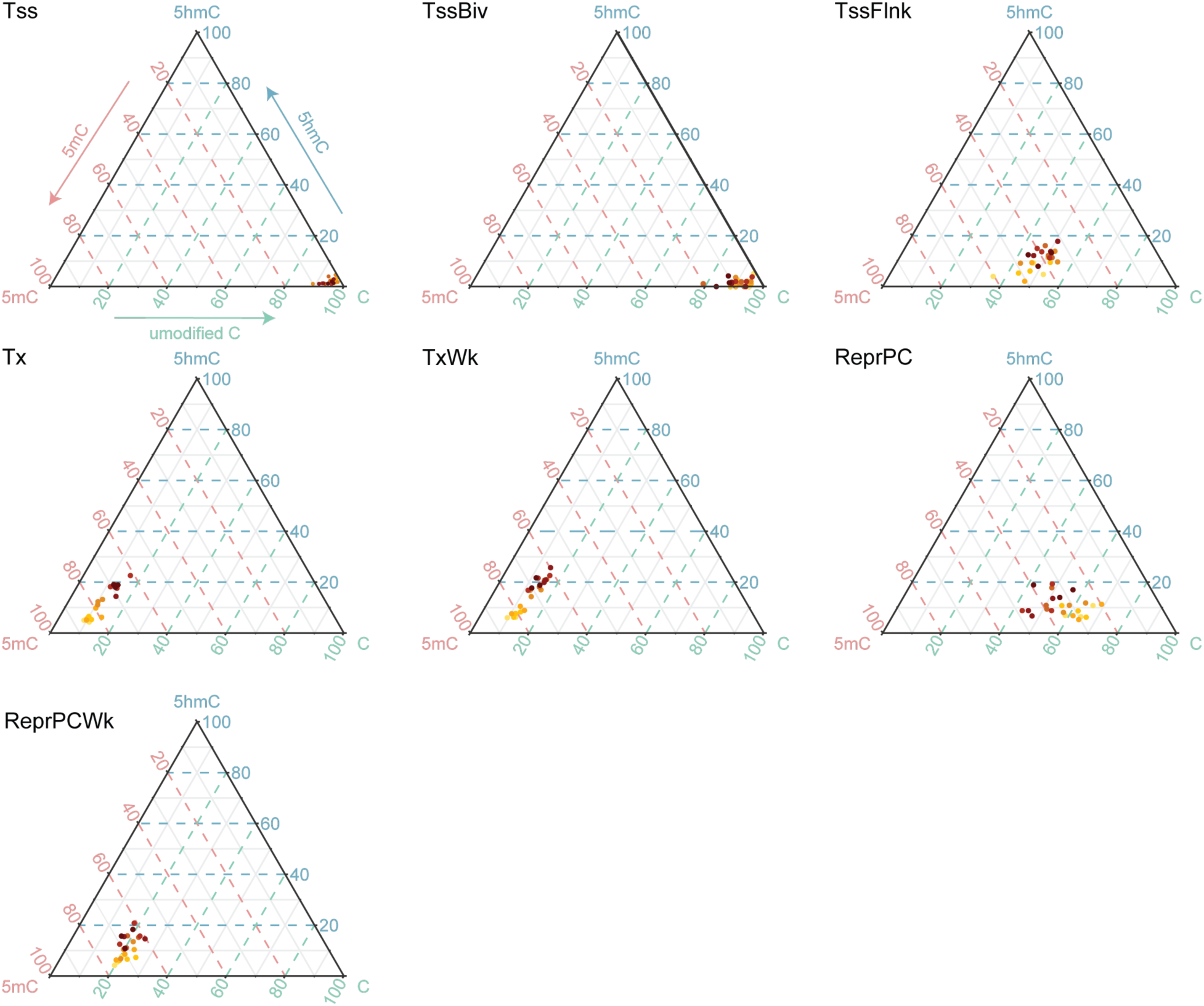
Ternary plots for change in DNA modifications by genomic element over development. Shown are ternary plots representing the levels of C, 5mC, and 5hmC measured for the whole genome or specific genomic elements. Each data point represents the measurement from a single biological replicate, with different time points represented by different colors.

**Figure S7.**
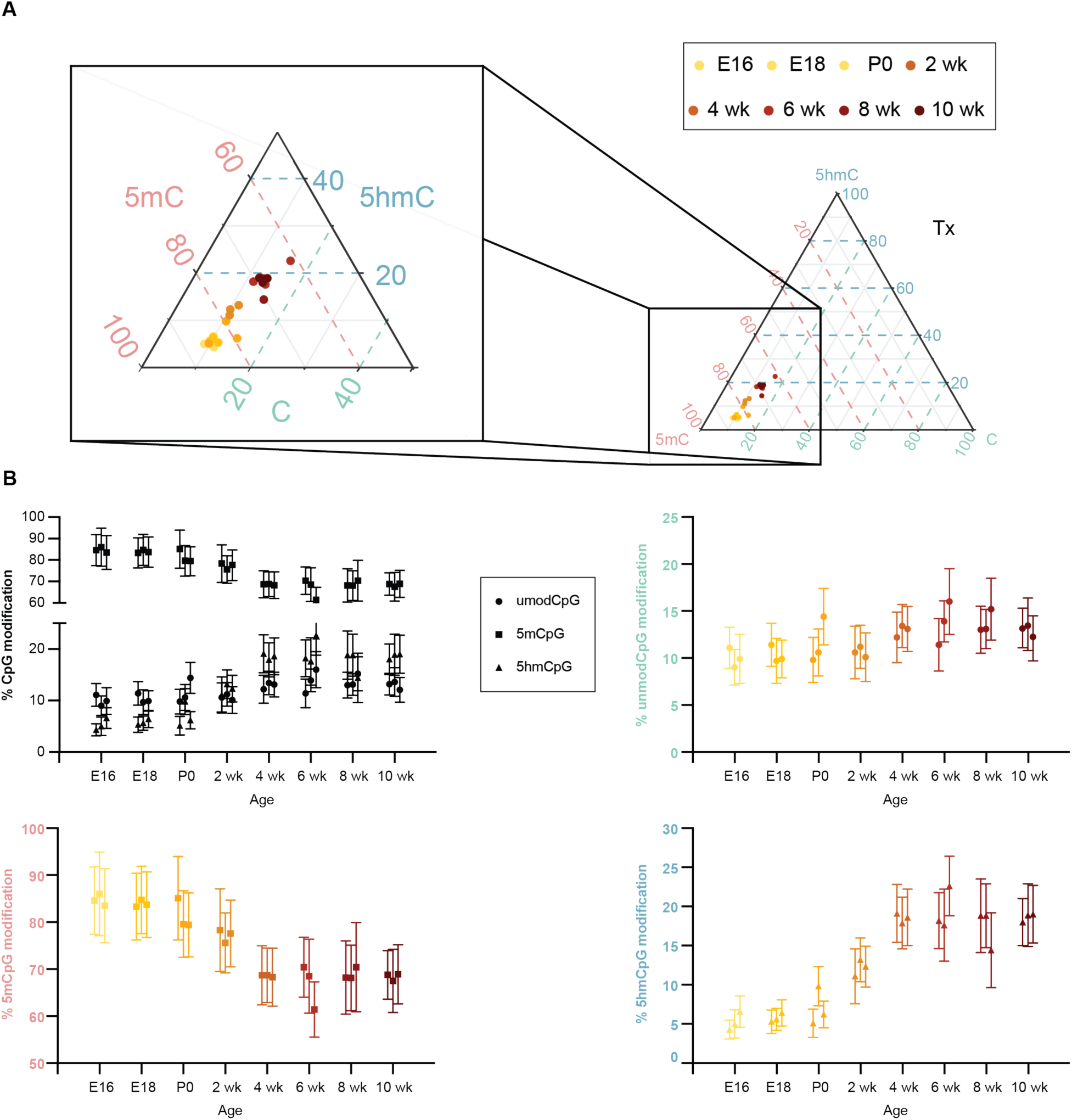
Total Analytical Error (TAE) for unmodified CpG, 5mCpG, and 5hmCpG measurements at transcribed elements across murine development. **A)** A ternary plot of the Tx chromatin state across the murine brain development, detected by Sparse BS/bACE-Seq, reveals that unmodified CpG levels remain stable while 5mCpG levels decrease and 5hmC levels increase over time. A zoom in of the relevant region of the ternary plot is shown. **B)** These dynamics are illustrated in scatter plots showing unmodified CpG, 5mCpG, and 5hmCpG plotted together (top left), unmodified CpG alone (top right), 5mCpG alone (bottom left), and 5hmCpG (bottom right). At each timepoint, three data points are shown, each representing an independent biological replicate. TAE bars derived from the TAE calculator are shown for each dataset, allowing for determination of trends that account for measurement error.

**Figure S8.**
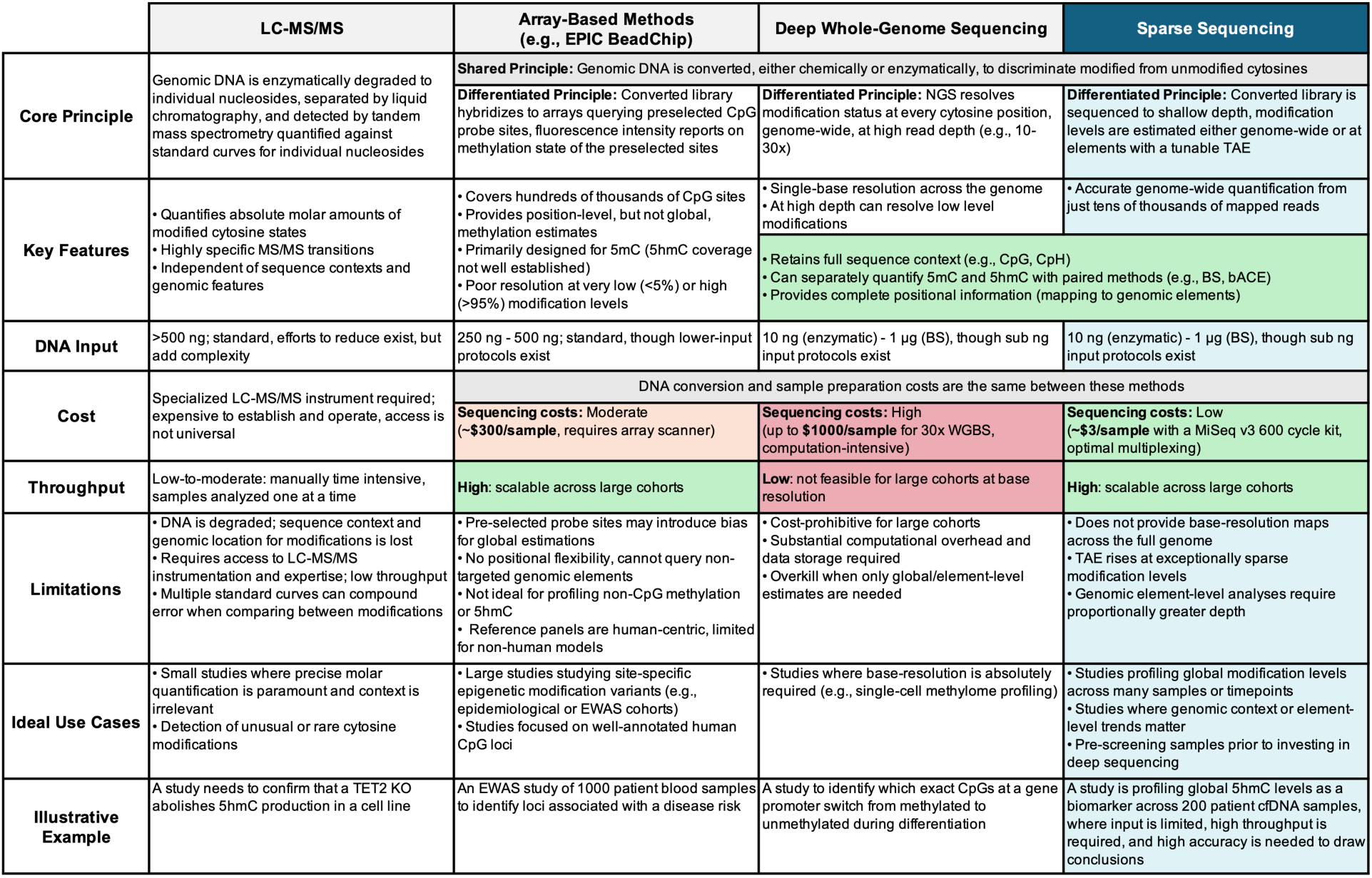
Comparison of methods for quantifying cytosine modifications. Four approaches for measuring unmodified cytosine, 5mC, and 5hmC are detailed and compared across key experimental and practical dimensions. Illustrative examples of use cases for each of these methods are also noted. Abbreviations: LC-MS/MS, liquid chromatography-tandem mass spectrometry; NGS, next-generation sequencing; TAE, total analytical error; bACE-Seq, bisulfite-assisted APOBEC-coupled epigenetic sequencing; WGBS, whole-genome bisulfite sequencing; EWAS, epigenome-wide association study; cfDNA, cell-free DNA.

### Supplementary Tables

**Table S1. Computational downsampling and derived Total Analytical Error (TAE)**

**Table S2. Total Analytical Error (TAE) as a function of genome coverage and % cytosine modification level**

## REFERENCES

1. Schubeler, D. (2015) Function and information content of DNA methylation. Nature, 517, 321–326.

2. Kohli, R.M. and Zhang, Y. (2013) TET enzymes, TDG and the dynamics of DNA demethylation. Nature, 502, 472–479.

3. Tahiliani, M., Koh, K.P., Shen, Y., et al. (2009) Conversion of 5-methylcytosine to 5-hydroxymethylcytosine in mammalian DNA by MLL partner TET1. Science, 324, 930–935.

4. Ito, S., Shen, L., Dai, Q., et al. (2011) Tet proteins can convert 5-methylcytosine to 5-formylcytosine and 5-carboxylcytosine. Science, 333, 1300–1303.

5. He, Y.F., Li, B.Z., Li, Z., et al. (2011) Tet-mediated formation of 5-carboxylcytosine and its excision by TDG in mammalian DNA. Science, 333, 1303–1307.

6. Pfeifer, G.P., Kadam, S. and Jin, S. (2013) 5-hydroxymethylcytosine and its potential roles in development and cancer. Epigenetics Chromatin, 6, 10–10.

7. Wagner, M., Steinbacher, J., Kraus, T.F., et al. (2015) Age-dependent levels of 5-methyl-, 5-hydroxymethyl-, and 5-formylcytosine in human and mouse brain tissues. Angew. Chem. Int. Ed Engl., 54, 12511–12514.

8. Globisch, D., Munzel, M., Muller, M., et al. (2010) Tissue distribution of 5-hydroxymethylcytosine and search for active demethylation intermediates. PLoS One, 5, e15367.

9. Booth, M.J., Raiber, E.A. and Balasubramanian, S. (2015) Chemical methods for decoding cytosine modifications in DNA. Chem. Rev., 115, 2240–2254.

10. Wang, T., Loo, C.E. and Kohli, R.M. (2021) Enzymatic approaches for profiling cytosine methylation and hydroxymethylation. Mol. Metab., 101314.

11. Amouroux, R., Nashun, B., Shirane, K., et al. (2016) De novo DNA methylation drives 5hmC accumulation in mouse zygotes. Nat. Cell Biol., 18, 225–233.

12. Vincent, J.J., Huang, Y., Chen, P.Y., et al. (2013) Stage-specific roles for tet1 and tet2 in DNA demethylation in primordial germ cells. Cell. Stem Cell., 12, 470–478.

13. Li, X., Wei, W., Zhao, Q.Y., et al. (2014) Neocortical Tet3-mediated accumulation of 5-hydroxymethylcytosine promotes rapid behavioral adaptation. Proc. Natl. Acad. Sci. U. S. A., 111, 7120–7125.

14. Stoyanova, E., Riad, M., Rao, A., et al. (2021) 5-hydroxymethylcytosine-mediated active demethylation is required for mammalian neuronal differentiation and function. Elife, 10, 10.7554/eLife.66973.

15. Gama-Sosa, M.A., Slagel, V.A., Trewyn, R.W., et al. (1983) The 5-methylcytosine content of DNA from human tumors. Nucleic Acids Res., 11, 6883–6894.

16. Chen, Z., Shi, X., Guo, L., et al. (2017) Decreased 5-hydroxymethylcytosine levels correlate with cancer progression and poor survival: A systematic review and meta-analysis. Oncotarget, 8, 1944–1952.

17. Chowdhury, B., Cho, I. and Irudayaraj, J. (2017) Technical advances in global DNA methylation analysis in human cancers. J. Biol. Eng., 11, 10–9. eCollection 2017.

18. Ellison, E.M., Abner, E.L. and Lovell, M.A. (2017) Multiregional analysis of global 5-methylcytosine and 5-hydroxymethylcytosine throughout the progression of alzheimer’s disease. J. Neurochem., 140, 383–394.

19. Morris-Blanco, K.C., Kim, T., Lopez, M.S., et al. (2019) Induction of DNA hydroxymethylation protects the brain after stroke. Stroke, 50, 2513–2521.

20. Irier, H.A. and Jin,P. (2012) Dynamics of DNA methylation in aging and alzheimer’s disease. DNA Cell Biol., 31 **Suppl 1**, 42.

21. Starczak, M., Gawronski, M., Olinski, R., et al. (2021) Quantification of DNA modifications using two-dimensional ultraperformance liquid chromatography tandem mass spectrometry (2D-UPLC-MS/MS). Methods Mol. Biol., 2198, 91–108.

22. Traube, F.R., Schiffers, S. and Carell, T. (2021) Quantification of DNA methylation and its oxidized derivatives using LC-MS. Methods Mol. Biol., 2272, 77–94.

23. Li, S. and Tollefsbol, T.O. (2021) DNA methylation methods: Global DNA methylation and methylomic analyses. Methods, 187, 28–43.

24. Taghizadeh, K., McFaline, J.L., Pang, B., et al. (2008) Quantification of DNA damage products resulting from deamination, oxidation and reaction with products of lipid peroxidation by liquid chromatography isotope dilution tandem mass spectrometry. Nat. Protoc., 3, 1287–1298.

25. Boulias, K. and Greer, E.L. (2021) Detection of DNA methylation in genomic DNA by UHPLC-MS/MS. Methods Mol. Biol., 2198, 79–90.

26. Dai, Y., Yuan, B. and Feng, Y. (2021) Quantification and mapping of DNA modifications. *RSC Chem*. Biol., 2, 1096–1114.

27. Darst, R.P., Pardo, C.E., Ai, L., et al. (2010) Bisulfite sequencing of DNA. *Curr. Protoc. Mol. Biol.*, **Chapter** 7, Unit 7.9.1–17.

28. Yu, M., Hon, G.C., Szulwach, K.E., et al. (2012) Tet-assisted bisulfite sequencing of 5-hydroxymethylcytosine. Nat. Protoc., 7, 2159–2170.

29. 29. De Borre, M. and Branco, M.R. (2021) Oxidative bisulfite sequencing: An experimental and computational protocol. Methods Mol. Biol., 2198, 333–348.

30. Caldwell, B.A., Liu, M.Y., Prasasya, R.D., et al. (2021) Functionally distinct roles for TET-oxidized 5-methylcytosine bases in somatic reprogramming to pluripotency. Molecular Cell, 81, 859–869.e8.

31. Fabyanic, E.B., Hu, P., Qiu, Q., et al. (2023) Joint single-cell profiling resolves 5mC and 5hmC and reveals their distinct gene regulatory effects. Nat. Biotechnol., 42, 960–974.

32. Schutsky, E.K., DeNizio, J.E., Hu, P., et al. (2018) Nondestructive, base-resolution sequencing of 5-hydroxymethylcytosine using a DNA deaminase. Nat. Biotech., 36, 1083–1090.

33. Vaisvila, R., Ponnaluri, V.K.C., Sun, Z., et al. (2021) Enzymatic methyl sequencing detects DNA methylation at single-base resolution from picograms of DNA. Genome Res., 31, 1280–1289.

34. Liu, Y., Siejka-Zielinska, P., Velikova, G., et al. (2019) Bisulfite-free direct detection of 5-methylcytosine and 5-hydroxymethylcytosine at base resolution. Nat. Biotechnol., 37, 424–429.

35. Wang, T., Fowler, J.M., Liu, L., et al. (2023) Direct enzymatic sequencing of 5-methylcytosine at single-base resolution. Nature Chem Bio, 19, 1004–1012.

36. Harris, R.A., Wang, T., Coarfa, C., et al. (2010) Comparison of sequencing-based methods to profile DNA methylation and identification of monoallelic epigenetic modifications. Nat. Biotechnol., 28, 1097–1105.

37. Weisenberger, D.J., Campan, M., Long, T.I., et al. (2005) Analysis of repetitive element DNA methylation by MethyLight. Nucleic Acids Res., 33, 6823–6836.

38. Yang, A.S., Estécio, M.R.H., Doshi, K., et al. (2004) A simple method for estimating global DNA methylation using bisulfite PCR of repetitive DNA elements. Nucleic Acids Res., 32, e38.

39. Kaur, D., Lee, S.M., Goldberg, D., et al. (2023) Comprehensive evaluation of the infinium human MethylationEPIC v2 BeadChip. Epigenetics Communications, 3, 6.

40. Ortega-Recalde, O., Peat, J.R., Bond, D.M., et al. (2021) Estimating global methylation and erasure using low-coverage whole-genome bisulfite sequencing (WGBS ). Methods Mol. Biol., 2272, 29–44.

41. 41. van der Velde, A., Fan, K., Tsuji, J., et al. (2021) Annotation of chromatin states in 66 complete mouse epigenomes during development. *Commun*. Biol., 4, 239–4.

42. Krueger, F. and Andrews, S.R. (2011) Bismark: A flexible aligner and methylation caller for bisulfite-seq applications. Bioinformatics, 27, 1571–1572.

43. Lister, R., Mukamel, E.A., Nery, J.R., et al. (2013) Global epigenomic reconfiguration during mammalian brain development. Science, 341, 1237905.

44. Wang, T., Luo, M., Berrios, K.N., et al. (2021) Bisulfite-free sequencing of 5-hydroxymethylcytosine with APOBEC-coupled epigenetic sequencing (ACE-seq). Methods Mol. Biol., 2198, 349–367.

45. Luo, C., Keown, C.L., Kurihara, L., et al. (2017) Single-cell methylomes identify neuronal subtypes and regulatory elements in mammalian cortex. Science, 357, 600–604.

46. Marchante-Gayón, J.M., Nicolás Carcelén, J., Potes Rodríguez, H., et al. (2024) Quantification of modified nucleotides and nucleosides by isotope dilution mass spectrometry. Mass Spectrom. Rev., 43, 998–1018.

47. Feng, J., Chang, H., Li, E., et al. (2005) Dynamic expression of de novo DNA methyltransferases Dnmt3a and Dnmt3b in the central nervous system. J. Neurosci. Res., 79, 734–746.

48. Chatterton, Z., Lamichhane, P., Ahmadi Rastegar, D., et al. (2023) Single-cell DNA methylation sequencing by combinatorial indexing and enzymatic DNA methylation conversion. Cell. Biosci., 13, 2–9.

49. Djirackor, L., Halldorsson, S., Niehusmann, P., et al. (2021) Intraoperative DNA methylation classification of brain tumors impacts neurosurgical strategy. Neurooncol. Adv., 3, vdab149.

50. Vermeulen, C., Pagès-Gallego, M., Kester, L., et al. (2023) Ultra-fast deep-learned CNS tumour classification during surgery. Nature, 622, 842–849.

